# Two interconnected patterning loops are required for body axis and head organizer formation in *Hydra*

**DOI:** 10.1101/2021.02.05.429954

**Authors:** Moritz Mercker, Anja Tursch, Frits Veerman, Alexey Kazarnikov, Stefanie Höger, Tobias Lengfeld, Suat Özbek, Thomas W Holstein, Anna Marciniak-Czochra

## Abstract

The formation of body axes and apical termini is crucial for animal development. In *Hydra*, nuclear *β*-catenin and Wnt3 play key roles and were previously thought to be part of a single mechanism for axis and head formation. This study challenges this view by combining mathematical modeling with experimental data. We show that *β*-catenin and Wnt3 patterning in Hydra operate at two different scales, requiring distinct inhibitory mechanisms. *β*-catenin, possibly interacting with other Wnts, co-ordinates axis formation, whereas Wnt3 is involved in small-scale head patterning. A double-loop reaction-diffusion model was developed, demonstrating the ability to describe patterns with divergent shapes, which single-loop models could not achieve. The previously proposed threshold mechanism for Wnt3 expression based on *β*-catenin prepatterns could not explain the data. Our results suggest a more complex patterning mechanism in other animals, where axis and head formation may not be controlled by a single process.

## INTRODUCTION

Understanding the mechanisms responsible for formation of the primary body axis, including the head, is a crucial issue in developmental biology (66). The process can be either based on maternally derived pre-patterns, such as in the fruit fly embryo (17), or based on a self-organized process, as demonstrated in various developmental systems ranging from simple metazoans (including *Hydra*) to complex mammalian organs (26; 64; 72; 86; 99). Moreover, even in vertebrate systems, recent data have revealed that spontaneous symmetry breaking occurs in the absence of extra-embryonic tissues (5; 84; 87). Consequently, self-organization may serve as a mechanism to ensure the robustness of developed structures under various intrinsic and extrinsic conditions.

Cnidarians provide a popular model system for studying general and transmissible developmental principles (21; 36; 43), and the freshwater polyp *Hydra* has been used to study development and regeneration for nearly 300 years. *Hydra* is used to systematically investigate axis and head formation under different natural and perturbed conditions. Head and axis patterning are constitutively active in adult polyps, to ensure maintenance of the body axis admits a continuous flow of self-renewing and differentiating cells (8; 9; 10). Moreover, the system regenerates ‘extremities’, such as the head and foot, as shown by Trembley in 1740 (83) and subsequently demonstrated in a range of tissue manipulation experiments (7; 61). Notably, *Hydra* can also regenerate entire polyps from dissociated and re-aggregated cells (26), providing a paradigm for *de novo* pattern formation (71; 80). Furthermore, *Hydra* tissue pieces can be transplanted from a donor to a host polyp by ‘grafting’, thus providing a model for mutual spatial interactions between tissue pieces from different body regions and different morphogenetic backgrounds (16; 48; 49; 73; 102). Body-axis and head formation also occur during budding, representing *Hydra*’s asexual form of reproduction, where buds evaginate and grow in the budding region of the mother polyp, eventually detaching as functional polyps (65). Finally, because *Hydra* also undergoes sexual reproduction, the mechanisms driving axis and head formation during embryogenesis can also be studied (19; 24; 52).

The mechanisms controlling the formation, regeneration, and maintenance of the body axis, including the head, remain an important issue in *Hydra* research (20; 28; 77; 94). Early grafting and regeneration experiments suggested the existence of so-called positional information, which informed cells about their relative position along the body axis, thus determining their functions (16; 26; 48; 49; 83). Recent studies identified several candidate molecules involved in these processes, including nuclear *β*-catenin (hereafter *β*-catenin) and HyWnt3 (hereafter Wnt3). These two molecules were identified as key players in head and axis formation in *Hydra* (15; 23; 34; 79), suggesting that canonical Wnt signaling plays an important role in the patterning process. Canonical Wnt signaling has also been intensively studied in a range of other model organisms, and Wnt activity, linked to translocation of *β-*catenin into the nucleus, was shown to be involved in the control of various developmental processes (reviewed in (4; 41)). These results led to the conclusion that the canonical Wnt signaling pathway constitutes the core structure of the self-organized patterning system in *Hydra* to simultaneously form the body axis and the head.

Such self-organization is usually described by systems based on a local self-activation coupled to a long-range inhibition (LALI) mechanism (42; 56; 90). The LALI principle is a generic concept that underlies several classes of pattern formation models, such as Turing systems of diffusing morphogens (85), cell-localized non-linear feedback models based on the coupling of diffusive and non-diffusive system components (51; 58) or mechano-chemical models (57; 59). Its most prominent implementation is the so-called activator-inhibitor model (27; 53; 54; 56). It is based on the Turing theory that describes symmetry breaking as a bifurcation from a spatially homogeneous state driven by contrasting diffusivity in a reaction-diffusion system of molecules. In line with the activator-inhibitor model, it has been suggested that Wnt3 and *β*-catenin form a self-activating loop via canonical Wnt signaling (36; 53; 54), leading to a concerted effort to identify the appropriate long-range inhibitor by experimental means (93; 103). The objective was to ascertain the minimal pattern formation system necessary for axis and head formation in *Hydra*, i.e. a molecular network that is sufficient to induce *de novo* patterning, based on homogeneous or randomly perturbed initial conditions without prepatterns.

Despite the significant progress that has been made since the publication of the activator-inhibitor model, an experimental identification of a chemical interaction network that can explain *de novo* pattern formation in *Hydra* remains elusive. This lack of success may be partly related to the growing evidence of the role of mechanical interactions in *Hydra* patterning (96). However, it is also possible that the models previously used to guide the *Hydra* research were inadequate. In this context, both Wnt3 and *β*-catenin have been used as interchangeable components of the central self-activation loop that ultimately leads to head and axis formation. Consequently, previous experiments have assumed a redundant phenotype when either Wnt3 or *β*-catenin is manipulated by genetic or pharmacological treatment (93; 98; 103). However, if this fundamental assumption is invalid and the molecules are controlled more independently in the patterning process, then future experiments may enable coherent insights into the overall process by identification of the respective patterning networks.

Further considerations require a clear distinction between the properties and roles of Wnt3 and *β*-catenin at two different levels. First, there is the ‘intrinsic function’, which refers to the induced biological processes, i.e. the intrinsic molecular properties and activity of a biological molecule, including its biochemical function, regulatory mechanisms - such as promoter control or constitutive activity - and direct role in cellular processes. Second, we consider ‘patterning function’ which refers to their role in network interactions underlying the self-organized pattern formation. In terms of ‘intrinsic function’ there are clear differences between the two molecules: The diffusible, extracellular protein Wnt3, for example, is strongly associated with the head organizer and thus with the formation of the head (34; 44). The role of the intracellular protein *β*-catenin is much more versatile: As a transcription factor, it is involved in the expression of various genes, but it is also a structural component of cell-cell contacts (97). However, these differences do not provide insight into their similarity with respect to their ‘patterning function’, i. e. whether the respective pattern formation is controlled by a joint long-term inhibitory mechanism.

Consequently, the study of ‘patterning function’ cannot only refer to the different phenotypes resulting from the manipulation of *β*-catenin or Wnt3, but has to consider specific scenarios concerning the pattern formation process, such as (1) spatial and temporal pattern overlap, e.g. if patterning of the two molecules is based on the same feedback-loop, the patterns should have a strongly related shape; (2) strength of coupling between both components, i.e. if they have a similar patterning function, expression levels of both molecules should distinctly depend on each other; (3) the question of whether and which patterns (head/ body axis) can develop if one of the molecules is inactivated (directly targeting the unique role and hierarchy of both molecules and patterning processes); and (4) the effect of the perturbed levels of both molecules on the levels of known inhibitors (if they have a similar patterning function, their effect on inhibitors should be qualitatively comparable).

In this paper, we assess these questions systematically by using mathematical models and analyses of previous and new experimental data. We explore the possibilities of a model based on a single coupling loop. To this end, we revisit the late Meinhardt model, integrating *β*-catenin and Wnt3 into a single patterning mechanism (54; 55). We further test such mechanism using new prototype models of a system comprising *β*-catenin, Wnt3 and a single long-range inhibitor. The discrepancy between model results and experimental observations, prompt us to extend the existing models to describe the nuclear dynamics of *β*- catenin and Wnt3 based on two patterning systems. These models, validated by previous and newly generated experimental data, suggest that *β*-catenin translocation (in concert with as yet unknown factors such as other Wnts) is involved in a large-scale body axis formation, while Wnt3 drives a separate small-scale patterning system responsible for organizer formation. The results of this study indicate that the control mechanisms may also be more complex in other animals, where canonical Wnt signaling has been assumed to be the main driver of body-axis (including head) formation (66). Given that Wnt/*β*-catenin- mediated pattern formation is fundamental to animal development, our data might also have implications for higher bilaterians, including humans.

## RESULTS

### Single- and double-loop models of Wnt3 and *β*-catenin patterning

To investigate the relationship between body axis and head patterning, we formulate two different mathematical models: a single-loop model based on a single pattern formation system and a new double-loop model coupling two patterning systems. Details of the mathematical setup are given in the Materials and Methods section. In the following, we focus on the introduction of the models, their biological justification and the model validation based on experimental data.

First, we formulate different hypothetical interaction networks extending the classical activator-inhibitor system to account for Wnt3 and *β*-catenin patterning in *Hydra*. We choose the activator-inhibitor model as a specific phenomenological description of the LALI (Local Activation Long-range Inhibition) principle. As mentioned above, the LALI principle might be realized in a different way in *Hydra*, e.g. involving mechanical inhibitors (96). However, as distinguishing between the specific long-range inhibitory mechanisms is not the scope of this work, we limit this study to the mathematical framework of reaction-diffusion equations. The link between Wnt3 and *β*-catenin is modeled following the canonical Wnt signaling and the mutual positive influence as demonstrated in various *Hydra* studies (see Supporting Information S1).

Previous works on expression patterns of *wnt3* vs. *β-catenin*, reveal that *wnt3* is expressed in small sharp spots compared to the rather large-scale patterns of *β-catenin* expression (15; 23; 34). The latter suggests that patterning of *β*-catenin, as a cell-localized molecule, develops in interaction with long-range mechanisms, e.g. diffusive molecules or mechanical tissue properties. Previously, this remarkable discrepancy in the size of the expression spot was modeled by a threshold-dependence assuming that Wnt3 production is locally enforced once a certain *β*-catenin threshold is exceeded (55). Resembling Wolpert’s seminal theory of positional information (101), the latter assumes a single-loop body-axis/head patterning system which is based on an activator-inhibitor model. The threshold-based function describes *wnt3* expression as a subset of the broader *β- catenin* expression domain (Figure 1). In interplay with a long-range inhibitor, this mechanism can explain both, self-organized pattern formation and the difference in expression spot size. Importantly, it also requires both molecules for the patterning process (cf., network A in Figure 1). A related network can be also constructed, where only one of both molecules is required for the patterning mechanism (cf., network B and C in Figure 1) and the other molecule is connected through canonical Wnt signaling and thus fulfills its intrinsic function aligned to the established pattern. In addition to these different single-loop models, we propose a double-loop system (cf., network D in Figure 1) which does not rely on any threshold mechanism but different expression spot sizes result from the intrinsic properties of two separate pattern formation loops, which are connected through canonical Wnt signaling.

**Figure 1.**
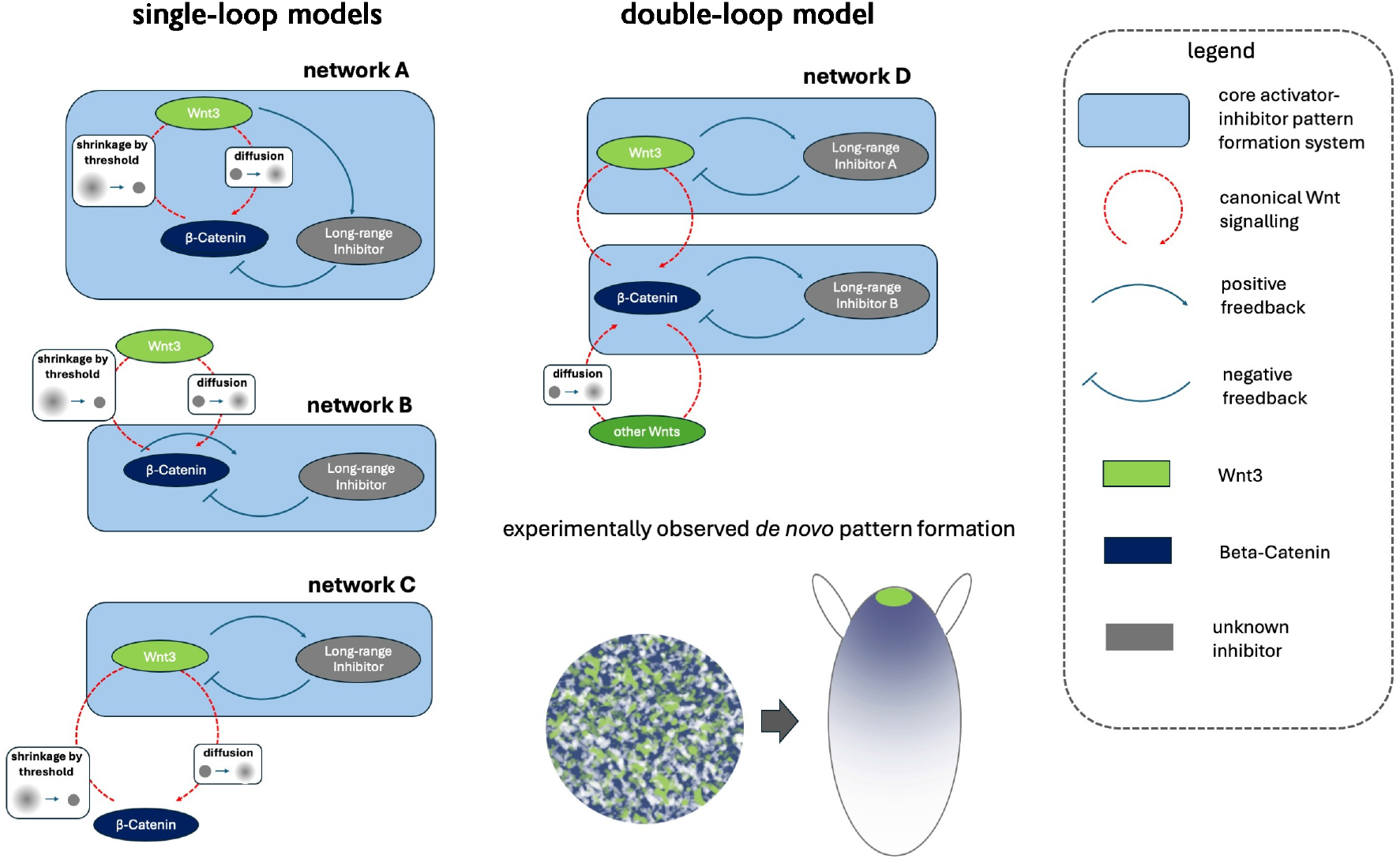
Four different possible network classes explaining *de novo* pattern formation of *β*-catenin and Wnt3 in *Hydra*. Networks comprise canonical Wnt signaling and a mechanism explaining experimentally observed difference in expression domain size between both components. Long-range inhibitors might be of molecular or of other nature (e. g., mechanical).

Networks A, B and C are based on a single pattern formation system and a threshold function-based dependence of Wnt3 on *β*-catenin. Since the simulations of the corresponding models show a qualitative similarity, in the following we focus on network B as a representative of the prototypical single-loop model. In this context, it is noteworthy to mention the proposal of a threshold-based model for Wnt3 and *β*-catenin by the late Meinhardt, in conjunction with numerous other molecular details and a hypothetical inhibitor (54; 55). However, systematic simulations of the model proposed by Meinhardt (2012, 2012a) show the emergence of irregular, growing-in-time spike patterns, and thus its inability to generate robust *wnt3* expression patterns (see Supplementary Information S3). This phenomenon can be elucidated through the analysis of reaction-diffusion-ODE systems with Turing instability. Such systems have been observed to exhibit irregular spikes for a specific choice of nonlinearities (32; 50; 70). This observation allows the rejection of the model.

In order to further explore the capacity of a single-loop model based on threshold mechanisms to reproduce the experimental data, we reduce the molecular complexity of the single-loop Meinhardt model and formulate a basic model coupling the threshold assumption of the dependence of *wnt3* expression on nuclear *β*-catenin concentrations with an activator-inhibitor model for *β*-catenin patterning (modeling network B).

As an alternative, we propose a model based on network D, denoted as a double-loop model, which is motivated by several experimental and theoretical considerations (cf., Discussion). The model assumes that *wnt3* expression patterns are regulated by a partially autonomous patterning system rather than being a mere readout of *β*-catenin activity. The novelty of the double-loop model is that it zooms out the head/body axis patterning system and explicitly distinguishes between two interconnected patterning systems that act differently in the apical region versus the body column. The model assumes that the body-scale patterning system is informed by nuclear *β*-catenin in interplay with long-range system components, such as other Wnts or Dkk1/2/4 (cf., Discussion section). It activates, among others, the production of Wnt3, which in turn is mediated by an additional, independent control loop providing a head-organizing function at the small scale (cf., Figure 1 network D). Specific interactions between model components, assumptions about model parameters, initial conditions, and information on the simulation code and software are provided in the Materials and Methods section.

The basic structure of both patterning systems (the *β*-catenin and Wnt3 submodels) in the double-loop model is similar to the single-loop model. Both systems are based on the activator-inhibitor scheme that exhibits Turing instability. The use of the activator-inhibitor schemes in the models do not necessarily reflect the molecular nature of the patterning process. Instead, they act as substitutes for more complex underlying processes and mechanisms that are not yet well understood at the molecular level. Notably, as stated above, these mechanisms fit into the general LALI concept and could by replaced by other symmetry breaking LALI scheme. To demonstrate it, we check that comparable simulation results are obtained when replacing the entire patterning units of the model by alternative mechanisms, i.e., replacing the activator-inhibitor loop by a mechano-chemical one (see Supplementary Information S4).

### Spatial structure of expression patterns

If two molecules are part of the same pattern formation loop, their interactions, e.g., through mutual activation or inhibition, are governed by shared reaction-diffusion dynamics, which dictate a common spatial wavelength as a result of a bifurcation from a selected simple eigenvalue. As a result, maxima of the pattern must align either in phase or antiphase. It is not possible for one system component to possess a significantly higher or lower number of maxima without compromising the established scale of the mechanism. This principle remains valid even in scenarios where the sizes of the expression domains differ considerably, such as in instances where they are governed by a threshold mechanism (as in the single-loop model). In order to address this observation in the two investigated types of models, a series of simulated experimental scenarios (related to elevated *β*-catenin- or Wnt3-levels) is conducted and subsequently compared with the relevant experimental data. In particular, we simulate five different scenarios with both models, namely: (1) the undisturbed scenario, (2) early and late stages after transient activation of *β*-catenin upon Alsterpaullone (ALP) treatment, (3) global genetic overexpression of either *β-catenin* or (4) *wnt3*, as well as (5) knockdown of a Wnt3 inhibitor (e.g., *sp5* (93)). The simulation of the single-loop (2 (a)) and the double-loop model (2 (b)) yield in majorly similar outcomes, but they also exhibit differences under several conditions. Compared to the single-loop model, only the double-loop model is able to reproduce multiple transient *wnt3*-expression spots as observed experimentally (15; 23; 93). Additionally, the single-loop model predicts Wnt3 spots of a larger size upon *β*-catenin and particularly *wnt3* overexpression, which is not detected in the double-loop model. Another striking difference is observed for elevated Wnt3 levels due to overexpression or *sp5* knockdown. Only the double-loop model predicts the formation of multiple heads and axes, respectively, which rather meets the experimental data. By contrast, the single-loop model has never generated more than one head or axis in this context. This is also why we neglect the single-loop model from our analyses of the pairwise distances presented in the next section, because an estimation of probability distribution is not possible.

To further confirm that both components are associated with patterns at different scales, we measure pairwise distances between *Hydra* heads in experiments and simulations based on the overexpression of *β-catenin* and *wnt3* (Figure 2 (c)-(f) as well as Supporting Information S2). The results demonstrate in both, experimental data (Shapiro-Wilk test: *W* = 0.80, *p* < 0.001, Figure 2 (d)) and simulations (Shapiro-Wilk test: *W* = 0.87, *p* < 0.001, Figure 2 (f)), that Wnt3 elevation (via *sp5*-knockdown) leads to a bimodal pattern of pairwise distances between heads, indicating that multiple heads cluster locally. Our *in silico* predictions (Figure 2 (b)) as well as previous experiments (93) reveal that these clusters appear close to the head and in the budding region, which are regions exhibiting high *β*-catenin (23; 34). Conversely, *β-catenin* overexpression leads to equally spaced heads, demonstrated via a unimodal pattern of pairwise distances (experiments: Shapiro-Wilk test: *W* = 0.96, *p* = 0.17, Figure 2 (c); simulations: Shapiro-Wilk test: *W* = 0.96, *p* = 0.81, Figure 2 (e)). This is in line with the phenotype of *β*-catenin overexpressing polyps, which show evenly spaced heads and feet along the axis (Figure 2 (h)). In summary, evaluations of simulations and experiments indicate that although Wnt3 and *β*-catenin are functionally intimately linked to direct patterning of axis/head structures, they do so on different spatial scales, which is only qualitatively reflected by the double-loop model.

**Figure 2.**
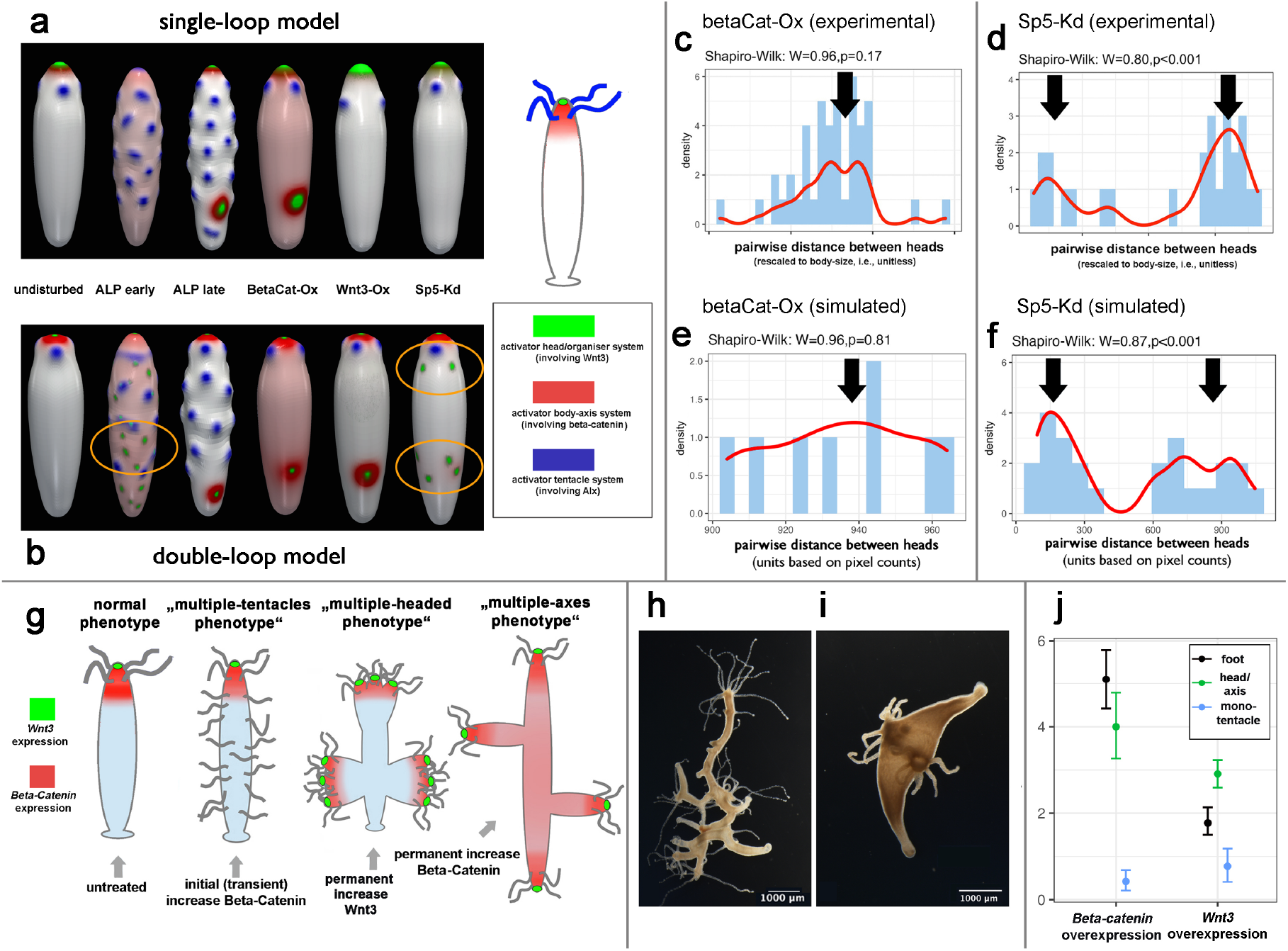
Virtual and experimental investigation of differences in spatial scales between *β*-catenin vs. Wnt3 patterns. (a)-(b): simulation snapshots of the single-loop model (a) vs. the double-loop model (b) for different scenarios (undisturbed, ALP = Alsterpaullone treatment, BetaCat-Ox = *β-catenin* overexpression, Wnt3-Ox = *wnt3* overexpression, Sp5-Kd = knockdown of the Wnt3-inhibitor (e.g., *sp5*)). Orange ovals indicate regions where the double-loop model reproduces experimentally observed multiple *wnt3*-expression spots. (c-f): Pairwise distances between heads in *β-catenin* overexpressing animals (c,e) vs. *wnt3* overexpressing animals (d,f) measured in experiments (c,f) vs. simulations of the double-loop model (e,f). (g) Scheme of different phenotypes resulting from different experimental conditions leading to *β*-catenin respectively Wnt3 increase; (h)-(i) darkfield images of phenotypes resulting from permanent *β*-catenin (h) vs. Wnt3 (i) increase; (j) statistical analysis of the number of feet, head and monotentacles for *β-catenin* vs. *wnt3* overexpression animals.

The implications of these pairwise distance analyses can be observed directly at the level of the induced phenotypes: An increase in Wnt3, following knockdown of Wnt3-specific inhibitors such as *sp5* (93) or *has-7* (103)), often results in the presence of a single body axis, accompanied by the development of multiple heads emanating from one *β*-catenin rich location (head, budding zone) (93; 103) (‘multi-headed phenotype’; Figure 2 (g-h)). The same occurs in *wnt3*-overexpressing animals, although the ectopic head locations are more variable (103). In contrast, a permanent and global increase in *β*-catenin results in animals with regular branching of new body axes (with each having a head at the apical end of the axis), with a body-scale distance between the newly formed axes (‘multiple axes phenotype’; Figure 2 (g-h)). Indeed, a statistical analysis of the number of heads/axes, feet and monotentacles (Figure2 (j)) demonstrates that for *wnt3*-overexpressing animals, the number of heads/axes is significantly higher than the number of feet (confidence intervals do not overlap) whereas in *β-catenin*-overexpressing polyps, confidence intervals strongly overlap. Notably, a transient increase of *β*-catenin induced by a high dose of ALP did not lead to the formation of multiple axes, but resulted in transient expression of multiple *wnt3* expression spots all over the body, followed by the formation of ectopic tentacles (15; 23) (‘tentacle phenotype’; Figure2 (g)).

### Strength of mutual coupling

If *β*-catenin and Wnt3 were part of a single common patterning loop, there would also be a direct correlation between the expression levels of the two molecules. In fact, it can be shown to be the case in all the single-loop models under consideration (cf., Figure 1, network A, B and C). To further investigate this point, we analyze the impact of *wnt3*-overexpression on *β-catenin* transcript levels and vice versa by comparing the results of qRT-PCR analysis to those of the single-loop and the double-loop model (Figure 3). Interestingly, high levels of *β*-catenin do not lead to significant changes in *wnt3* transcript *in vivo* (a) and *vice versa* (b). However, models of too strong overexpression lead either to numerical instabilities or to a complete loss of pattern formation. Thus, for each model, the maximum possible level of overexpression has to be set, which limits the comparability of simulations and experiments. In both models, the expression levels of both components appear even less decoupled than in experiments (Figure 3 (d)-(f)). Only assuming the double-loop model and concerning overexpression of *wnt3* allows a simulation of the strong virtual overexpression, the levels of which are one order of magnitude higher than those of *β-catenin* and thus comparable to the experimental result. Here, a distinct decoupling (similar to experiments) is observed (Figure 3 (f)), i.e., the relative increase of *wnt3* transcript is far beyond the resulting relative increase of *β-catenin* transcript.

**Figure 3.**
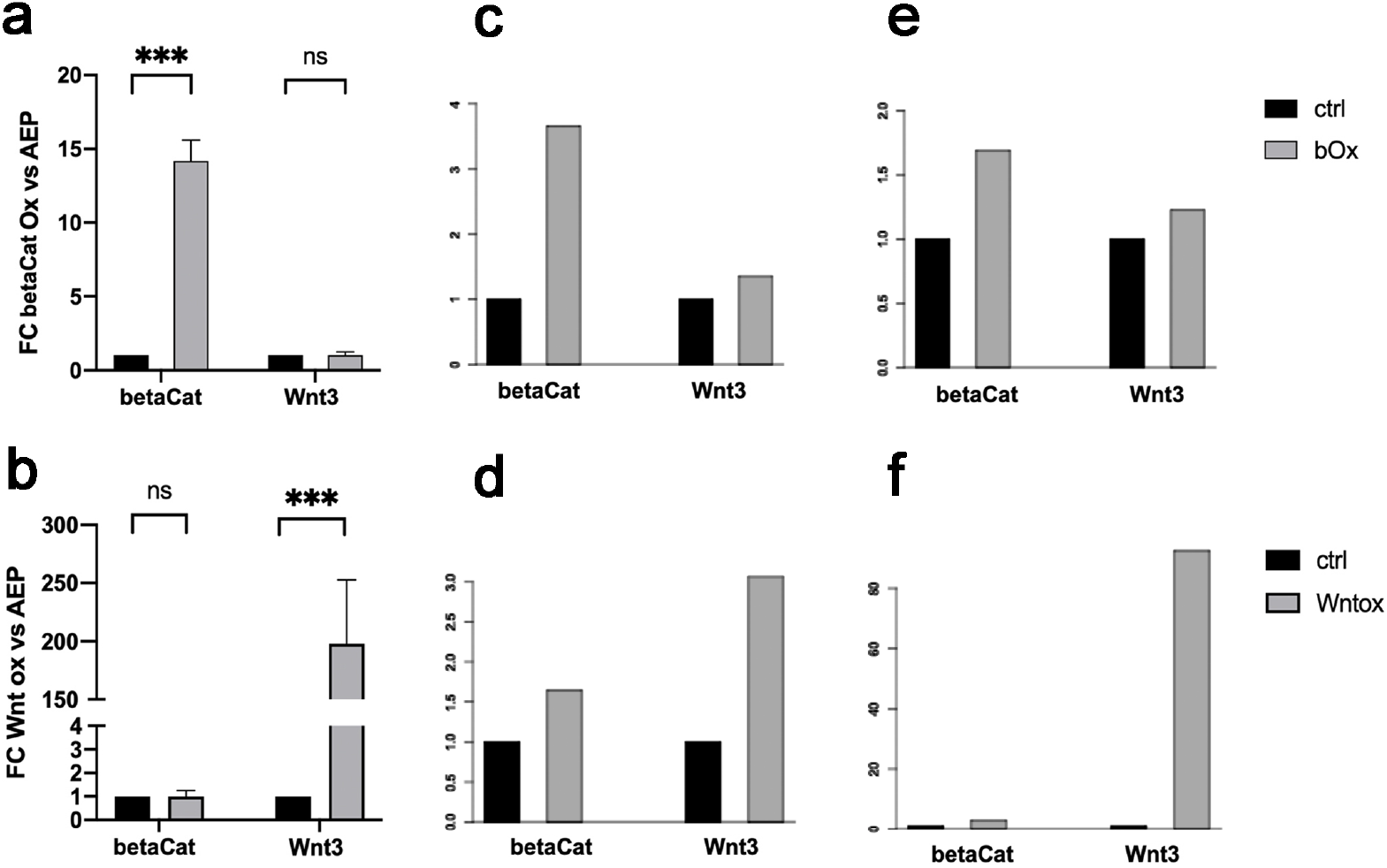
Investigation of the strength of mutual coupling between Wnt3 and *β*-catenin. Experiments (qRT-PCR analysis: (a-b)) and simulations of the single-loop model (integrated quantities: (c-d)) and the double-loop model (integrated quantities: (e-f)). (a),(c),(e) investigate overexpression of *β-catenin* (bOx); (b),(d),(f) investigate overexpression of *Wnt3* (Wntox). Levels are normalized to those of the control/wildtype scenario.

### Analysis of patterning function via pharmacological and genetic knockdown

It has already been shown that Wnt3 is necessary for head and organizer formation (e.g., (88)). However, it is unclear whether axis formation is an independent patterning process and whether Wnt3 or other Wnts are necessary for this. To gain further insight, we performed genetic knockdown and pharmacological inhibition experiments to investigate the effect of symmetry breaking and axis formation by determining the sphericity of reaggregates at day 6 after dissociation. With respect to the geometric measure, the interpretation is unambiguous; that is, the greater the spherical character of the aggregate (sphericity value approaching 1), the less probable is its recognition as a developed body axis. Conversely, the more developed the body axis, the smaller this value becomes. In later stages in fully functional animals, the shrinkage of this value is exacerbated by the elongation/growth of the axis. The corresponding results are shown in Figure 4 (exemplary phenoptypes: (a-b); statistical analysis: (c)). The control groups treated with either DMSO or electroporated with siGFP, show the lowest sphericity values indicative of proper axis development (Figure 4 (a-c)). In contrast, reaggregates silenced in *wnt3* exhibit defects in head patterning. However, axis formation is still detectable phenotypically, which is in line with the sphericity value, which was higher than in the control specimen but below the weighted average (Figure 4 (c)). In the event of canonical Wnt signalling being abolished by the knockdown of *wntless*, a cargo receptor required for general Wnt secretion, the majority of the reaggregates are unable to undergo both patterning and axis development (Figure 4 (a,c)). A similar result is obtained with reagregates treated with LGK-974, which blocks porcupine and thus general Wnt secretion and function (Figure 4 (b,c)). Conversely, exposure to iCRT-14, an inhibitor of *β*-catenin’s transcriptional activity, yield in similar results obtained upon inhibition of Wnt secretion (Figure 4 (b,c)). A relatively low concentration of iCRT has been used to ensure viability of the reaggregates, which most probably reduced the efficacy.

**Figure 4.**
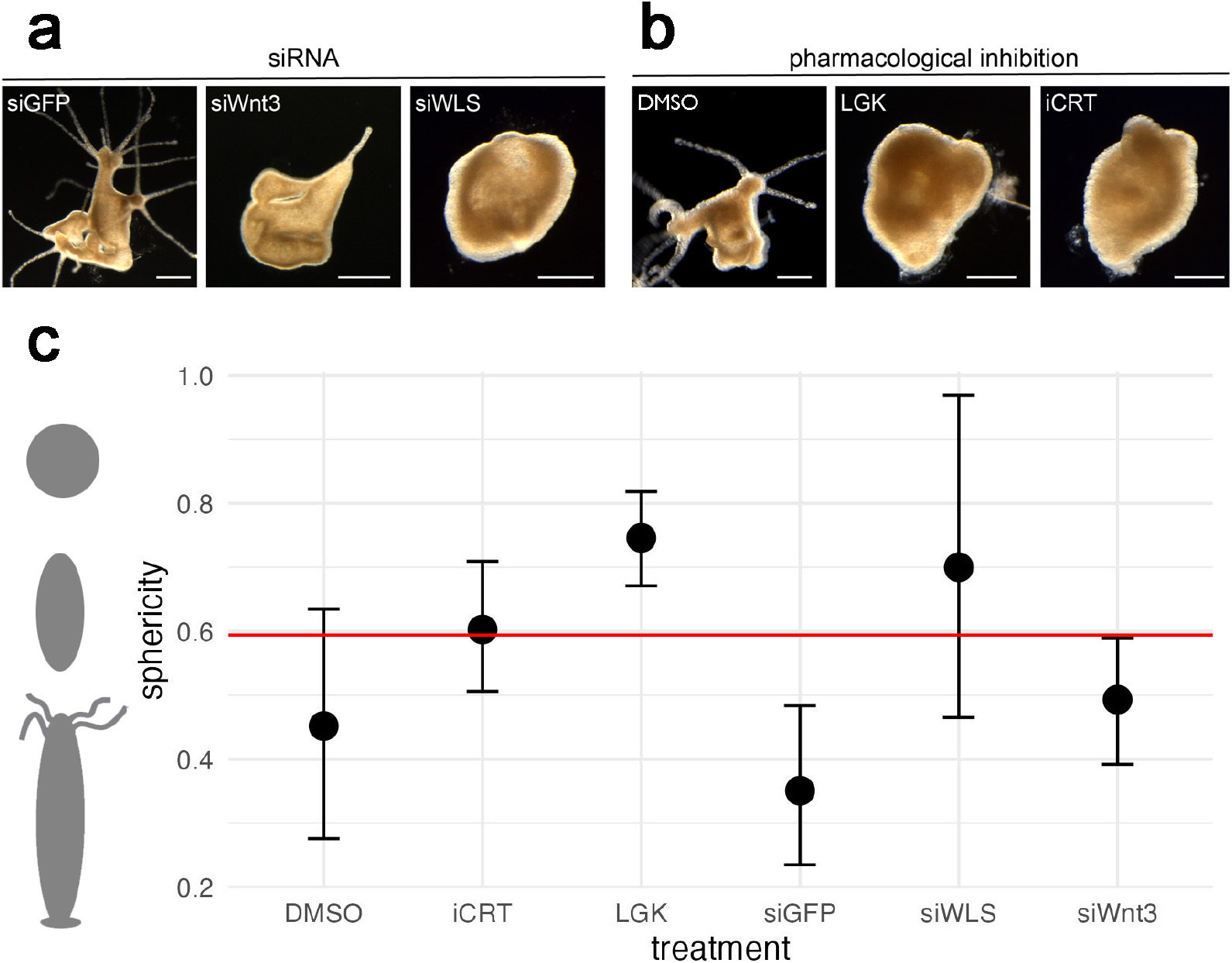
Investigating how different genetic (a) and pharmacological (b) knockdowns related to *β*-catenin, Wnt3 and Wnts in general influence the ability to establish a body axis in *Hydra* aggregates – the latter quantified by the average sphericity related to each treatment (c). (a)-(b): darkfield microscopical examples of different treatments (scale bars: 500 *μ*m), (c) statistical analysis of sphericity. DMSO = control group in pharmacological treatment, iCRT = pharmacological *β*-catenin inhibition, LGK = pharmacological inhibition of Wnt secretion, siGFP = control group in genetic knockdown, siWLS = genetic knockdown of *wntless* and siWnt3 = genetic knockdown of *wnt3*. Black dots represent average sphericity values, error bars 95% confidence intervals based on bootstrap and the red dotted line the weighted average sphericity value across all experiments.

### Interplay with other Wnts and known inhibitors

If Wnt3 and *β*-catenin contribute to the same activator complex of a single-loop patterning network, it is reasonable to assume that overexpression of either components has a qualitatively similar effect on expression levels of known Wnt antagonists. To address this, qRT-PCR analysis has been performed to check the relative expression levels of several Wnt genes as well as known antagonists of Wnt3 (Figure 5), (31; 93; 103). While Wnt 7, Wnt8 and Wnt9/10c do not respond to elevated Wnt3 or *β*-catenin levels, Wnt1 shows a down-regulation exclusively in *β-catenin* overexpressing animals. By contrast, Wnt2 is reduced only in transgenic Wnt3 polyps while it is elevated in *β-catenin* overexpressing animals, and Wnt11 levels are reduced in either strain used. With regard to the antagonists, HAS-7 was slightly up-regulated in *β-catenin* overexpressing polyps and strongly elevated in strains overexpressing *wnt3*. By contrast, Sp5 and Dkk1/2/4 showed a down-regulation upon increased Wnt3 levels, while high levels of *β*-catenin yielded in an opposing expression level for Dkk124. These data indicate that although *β*-catenin and Wnt3 are engaged in the same pathway within a positive feedback loop, their effects on a molecular level, are not mutually interchangeable, but partially distinct.

**Figure 5.**
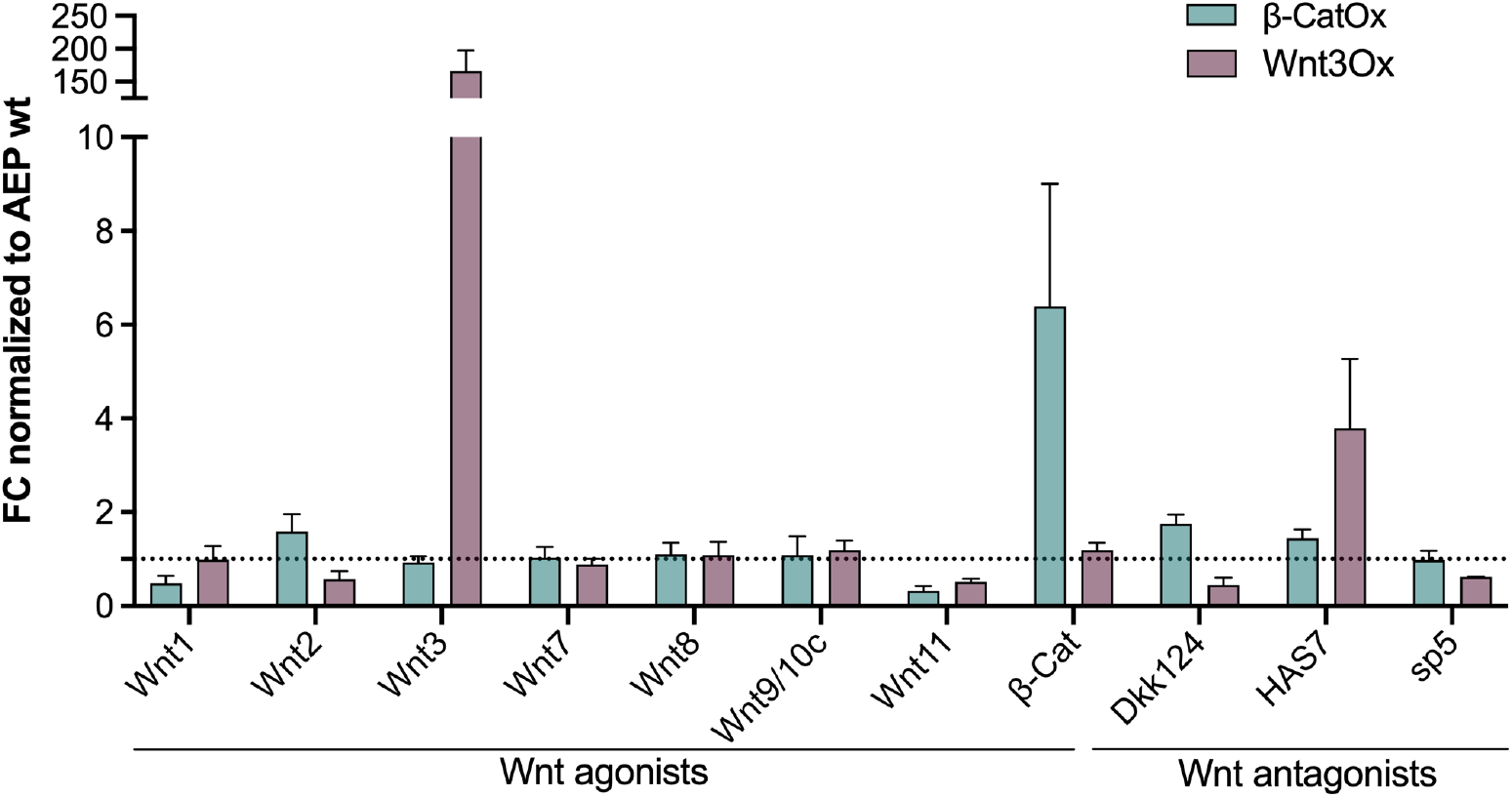
qRT-PCR analysis of Wnt agonists and antagonists in *β-catenin* (*β*-CatOx) and *wnt3* (Wnt3Ox) overexpressing animals. Values were normalized to ef1a and the wildtype animals served as a reference to obtain fold change expression. All samples were run in technical and biological triplicates. Bars represent mean values with standard deviation.

## DISCUSSION

The present work focuses on the ‘patterning function’ of Wnt3 and *β*-catenin molecules, with the aim of determining whether both molecules are part of a single patterning loop that generates head and axis simultaneously, or there are distinct regulatory mechanisms responsible for different spatial scaling of the observed patterns. The existence of two partially independent systems that are nevertheless still coupled through the canonical Wnt signaling is compelling from a developmental point of view, because it guarantees the robust formation and size of the Wnt3-associated organizer. Should the *wnt3* expression be contingent on the critical thresholds of *β*-catenin as postulated in (54; 55), then the manifestation and size of the organizer would be susceptible to spatiotemporal variations in *β*-catenin levels. Conversely, a high level of robustness would be achieved by two redundant patterning mechanisms acting at different spatial scales. This is a well-known principle in various biological contexts (95). Moreover, earlier studies on regeneration have demonstrated that the positioning of head versus foot formation is contingent on relative, as opposed to absolute, levels of the head-activation gradient (7; 8; 61). This observation renders a threshold-based mechanism even less probable.

In order to further investigate these hypotheses, two different models were simulated and contextualized with a variety of previous and new experimental data. The single-loop model represents the prevailing idea that both molecules act in a joint patterning loop (having canonical Wnt signaling as its core structure) and that head and axis formation thus occurs synchronously. This model needs however a threshold-based mechanism to explain the differences in expression domain size between *β-catenin* and *wnt3*. In contrast, the double-loop model is based on the assumption that the distribution of each molecule is controlled by a separate patterning process, coupled via canonical Wnt signaling.

A direct comparison of the different simulated scenarios reveals that both models adequately describe the steady-state scenario and can reproduce spontaneous head development (Fig. 2 (b) and (d) – undisturbed scenario). In particular, we observe a gradient-like pattern of *β-catenin* expression, with a maximum at the head, a small, sharp spot of *wnt3* expression at the tip of the hypostome, and a ring of tentacles beneath the head. Here, as in all other simulations based on the single-loop model, Wnt3 spots appear to be larger compared to the results of the double-loop model. Indeed, smaller spots could be obtained by applying a higher *β*-catenin threshold for *wnt3* expression in the single-loop model. However, in such cases, the appearance of Wnt3 is observed to be considerably later than that which has been experimentally determined. This discrepancy can be attributed to the patterning dynamics of the large-scale *β*-catenin system, which has been shown to generate gradients with a higher rate of speed in comparison to the relatively gradual increase in maximum levels. Furthermore, incorporating stochasticity into the levels of *β*-catenin would result in the occurrence of *wnt3* expression spots being contingent on the setting of thresholds that are not too narrow. Both of these illustrate the lack of robustness of the *wnt3* expression and the dependence of spot size on the threshold mechanism.

Also the various simulations of the manipulated scenarios in comparison with previous experimental data favor the double-loop model: Simulations with initial conditions corresponding to ALP treatment reproduced the experimentally observed transient multiple *wnt3* expression spots (e.g., (15; 23)) in early stages after ALP application in the double-loop model, but not in the one-loop model (Fig. 2 (a),(b) – cf., orange ovals). The same qualitative difference between both models holds when comparing simulations including knockdown of the Wnt3 inhibitor: here, the double-loop model generates multiple Wnt3 spots in both, the head region and the budding zone (Fig. 2 (b) orange ovals), in accordance with observations of the multiple-headed phenotype following *sp5* knockdown (Ref. (93)). However, the one-loop model was not able to reproduce these patterns.

Interestingly, in the late stages of simulations based on the double-loop model, ectopic Wnt3 spots vanished (not shown), whereas in real experimental systems, they developed into ectopic heads (93). Nevertheless, the ectopic Wnt3 spots persisted significantly longer in the simulated *sp5* knockdown model compared with simulations including ALP treatment. We concluded that a certain morphogen-synthesis time is required in the real system, sufficient to initiate a local cell differentiation process. The latter leads to ‘fixation’ of the organizer and suggests that a transient expression pattern is sufficient to control the process. These effects were not incorporated in our present and previous models, which is why the *wnt3* expression spots finally vanished following simulated *sp5* knockdown treatment.

In addition, also the simulations with respect to *wnt3* overexpression lead to more realistic results in the double-loop model: Here, the simulated pattern strongly resembles the *β-catenin*-overexpression phenotype in terms of the establishment of a secondary body axis, in accordance with the experimental observations (103) and in contrast to the single-loop model (Figure 2 (a-b)). However, both models do not reflect the experimental observation of ectopic tentacles, possibly because they do not consider the interaction of the tentacle system with the other systems in detail.

The observation that simulated *wnt3* overexpression in the double-loop model does not lead to a ‘multiple-headed phenotype’ as observed for *sp5* knockdown, but rather to a ‘multiple-axes phenotype’ (Fig. 2 (g)), may be initially surprising. However, coupling between the *β*-catenin-driven large-scale system and the Wnt3-driven small-scale system (due to canonical Wnt signaling) makes the situation more complex and less easy to predict. Notably, *wnt3* overexpression also activates *β*-catenin, which together leads to the formation of additional body axes in regions with already relative high *β*-catenin levels, e.g., the budding zone. In fact, the *wnt3* overexpression phenotypes observed experimentally are more variable and may show both ectopic axes and multiple heads (103). Overall, this suggests that the multiple head and multiple axis phenotypes cannot be used to conclusively assign manipulated molecules to one of the two systems, but rather represent the ends of a scale of possible phenotypes that may be difficult to interpret.

The presented work analyzed pairwise distances between heads in *β-catenin* overexpressing polyps vs. those where Wnt3 levels are elevated (via the knockdown of the Wnt3 inhibitor *sp5* – Figure 2 (c-f)) further support the observation that both molecules, *β*-catenin and Wnt3, are related to patterns acting at different scales: in both, experiments and simulations of the double-loop model, elevated *β*-catenin leads to regularly spaced additional body axes (the distribution of distances does significantly deviate from the normal distribution) whereas animals with elevated Wnt3 levels show distinct bimodal patterns, representing several heads/*wnt3* expression spots clustered in regions with higher levels of *β*-catenin, such as the apical end or the budding region. This mismatch clearly contradicts the assumption that both molecules are part of the same pattern formation system – since shared reaction-diffusion dynamics would dictate a common spatial wavelength. This is additionally supported by previous ALP experiments showing that high, body-wide, constant levels of *β-catenin* transcripts are not followed by *wnt3*, which was instead expressed in a complex arrangement of multiple transient organiser-sized spots all over the body (15; 23). This observation of scale-mismatch is also reflected at the phenotypic level, where Wnt3 elevation leads to multiple heads where only one head should arise (‘multiple head phenotype’ – Figure 2 (g), (j)), whereas *β-catenin* overexpression leads to multiple axes that resemble ladder structure (‘multiple axes phenotype’ – Figure 2 (g), (h)). This visual impression is again supported by the fact that Wnt3 elevated animals show significantly more heads than feet, whereas this difference is not significant in *β*-catenin elevated animals (Figure 2 (j)). Differences in total numbers of head/axis in both treatments in 2 (j) are however related to differences in total polyp length which is not further considered in our study. The observation that a transient increase in *β*-catenin levels (15; 23) (‘tentacle phenotype’; Figure2 (g)) or, alternatively, a moderate *β*-catenin increase (88) leads to ectopic tentacle formation in regions with elevated *β*-catenin levels might indicate a failure of the establishment of permanent *wnt3* expression centers.

The activities of Wnt3 vs. *β*-catenin differ not only in terms of their spatial scales, but also in their dynamics, further supporting the assumption that both molecules are integrated into different pattern formation networks. With the exception of injury-induced Wnt3 signaling (94), experiments on intact polyps suggested that *wnt3* was expressed at later stages than *β-catenin: Wnt3* expression was distinctly delayed compared with *β-catenin* during embryogenesis (19) and during budding, its expression commenced with the appearance of the first bud protrusions (34) whereas *β-catenin* expression is already evident prior to the first visible deformations (37; 97). Indeed, inhibition of *β-catenin* prevented bud formation, including the expression of several bud- and organizer-specific genes (97). However, caution is advised with regard to these temporal interpretations, since, on the one hand, analyses can be confounded with *β*-catenin activity related to its structural tasks and not so its signaling activity. On the other hand, Wnt3 may have relevant effects on pattern formation in much smaller quantities than *β*-catenin, which could lead to a supposedly later detection for reasons of test sensitivity. In summary, however, the results of our and previous spatial and temporal analysis strongly suggest that the patterning activities of both molecules are controlled by different mechanisms acting at different spatio-temporal scales, with *β-catenin* frequently preceding *wnt3* expression.

Our experimental analysis of the mutual coupling of beta-catenin and Wnt3 (3) shows that the expression of both components is almost completely decoupled, which is particularly surprising in the context of canonical Wnt signalling. This high degree of uncoupling can only be reproduced by the double-loop model with regard to the *wnt3* overexpression scenario, suggesting that in *Hydra* there are efficient control mechanisms, in particular for *β*-catenin-dependent *wnt3* expression, which are not yet reflected in the models. Indeed, a tight regulation of Wnt3 levels in *Hydra* is achieved through a combination of autoregulatory and repressive transcriptional mechanisms (93), cis-regulatory elements that confine expression to the head organizer (35; 63), extracellular antagonists like Dkk (31), and proteolytic degradation by the metalloprotease HAS-7 to prevent overproduction (103). The fact that in our simulations, the strength of coupling does not scale with the number of ectopic axes/heads (cf., Figure 2 (a-b)) but is more moderate, is most probably due to the fact that in the context of pattern formation, the different components are gradually distributed along the body axis and therefore represent a non-negligible quantity even outside their maxima. However, these coupling experiments and simulations have to be interpreted with care: apart from the missing additional Wnt3 level regulation mechanisms in our models (cf., above), in addition, our experimental results of a qRT-PCR to measure the signalling activity of nuclear *β*-catenin may be obscured to a greater extent by the structural activity of *β*-catenin. Finally, simulated overexpression strength between experiments and simulations strongly differ due to technical constraints (cf., Results section) which makes them comparable only to a limited extent.

On the function level, we finally proved via genetic and pharmacological knockdown of different components in *Hydra* aggregates that indeed, the inhibition of *β*-catenin (or alternatively all canonical Wnts) effectively prevents the formation of the body axis and aggregates remained spherical (showing the 3 highest sphericity values – Figure 4 (b), (c), (e)). This suggests, that body axis establishment requires the interplay between *β*-catenin and canonical Wnts, as previously assumed. The two control groups, in contrast, showed the lowest sphericity values (Figure 4 (a), (e)), which was expected as it reflects the establishment of the body-axis and subsequent outgrowth. *wnt3* knockdown, finally leads to slightly higher sphericity values that in the control groups (Figure 4 (d), (e)) but still significantly below the weighted average across all experiments. We conclude from this that indeed a body axis is established, further outgrowth is however avoided due to the lack of head development (88). Taken together, these results further support the assumption that *Hydra* axis vs. head formation are two partially decoupled pattern formation processes, where Wnt3 is responsible for organizer/head formation, but axis formation is driven by the interplay of *β*-catenin with other canonical Wnts. Indeed, also previous results demonstrate that axis formation (indicated by foot development) may take place even when suppressing Wnt3 activity by a lack of osmotically driven stretch (18) and in turn, a recent work on cnidarian embryos demonstrates that Wnt3 has a *β*-catenin independent role during patterning (89) – in agreement with our data indicating that Wnt3 is regulated by a patterning mechanism (including HAS-7) not directly regulated by *β*-catenin (cf., below).

Finally, we demonstrated that the influence of *β-catenin* vs. *wnt3* overexpression on three different Wnt antagonists (Dkk1/2/4, HAS-7 and Sp5) differs qualitatively and significantly in all cases (Figure 5): Sp5 and HAS-7 levels do not react distinctly on *β-catenin* overexpression but antagonistically on *wnt3* overexpression, fitting well with the previous assumption that these molecules are specific inhibitors of Wnt3 (93; 103). The distinct antagonistic reaction, however, suggests that their role is more complex than just representing the inhibitor molecule in an activator-inhibitor scheme – in accordance with the observation that both molecules partially contradict the properties as expected for the classical inhibitor from the theoretical point of view (93; 103). For this reason, both molecules could instead represent mechanisms that ensure that Wnt3 levels are robustly maintained at specific levels and the main pattern formation/inhibitory mechanisms might realized in a different, e.g., mechanochemical way (98). The distinct upregulation of HAS-7 based on Wnt3 suggests that HAS-7 functions as a specific Wnt3 inhibitor whose expression is not directly regulated by *β*-catenin, in line with the observation that it is not TCF-dependent (103) – and in contrast to sp5, being part of the *β*-catenin/TCF loop (93).

In contrast, Dkk1/2/4 distinctly (and antagonistically) reacts to both, *β-catenin* and *wnt3* overexpression. This might again suggest that Dkk1/2/4 is involved in *Hydra* body axis rather than organizer formation (most probably involving other Wnts – cf., Figure 4) as proposed by (58). Which further Wnts are involved in head vs. axis formation remains elusive. The analysis of the effect of Wnt3 vs. *β*-catenin overexpression on the levels of other canonical Wnts shows however a very heterogeneous picture in this regard, ranging from no reaction to a uniform reaction to an antagonistic reaction (Figure 5). This suggests that the tasks of Wnts in the context of head and axis formation and other systems (such as foot, source density and tentacles) is diverse and differentiated, which sheds a little more light on the question of why this diversity of Wnts (‘Wnt code’) is required (30).

In addition to the above-mentioned (and analysed) candidates for possible inhibitors of the body axis vs. organiser pattern formation system, further molecular and non-molecular candidates can be considered. For the small-scale (i.e., head-organiser-related) activator–inhibitor pattern formation system, in addition to HAS-7 and Sp5, Notum, another Wnt antagonist, is a carboxylesterase that requires glypicans to suppress Wnt signaling by depalmitoleoylation of Wnt proteins (38; 39). In the context of the body-axis system, one of different possible additional molecular candidates for the inhibitor complex is Naked cuticle (Nkd) which was shown to be expressed in broad, gradient-like patterns along the body axis resembling *β-catenin*/*tcf* expression after ALP-treatment (cf., Supporting Information S5). In addition, Nkd binds to Dishevelled (Dvl) family proteins preventing the translocation of *β*-catenin into the nucleus (69; 100) and in ALP experiments, it shows a distribution rather related to *β*-catenin than spotty patterns (as for Wnt3 and other Wnts – Supporting Information S5). However, due to its intracellular nature, the long-range inhibition cannot be explained by this protein alone but requires additional interactions, which can be also of a mechano-chemical, cellular or bio-electrical character (12; 18; 45; 59; 76) – as it could be the case for the Wnt3 pattern formation system (98). An overview of possible molecular candidates of the double-loop model is given in Figure 6.

**Figure 6.**
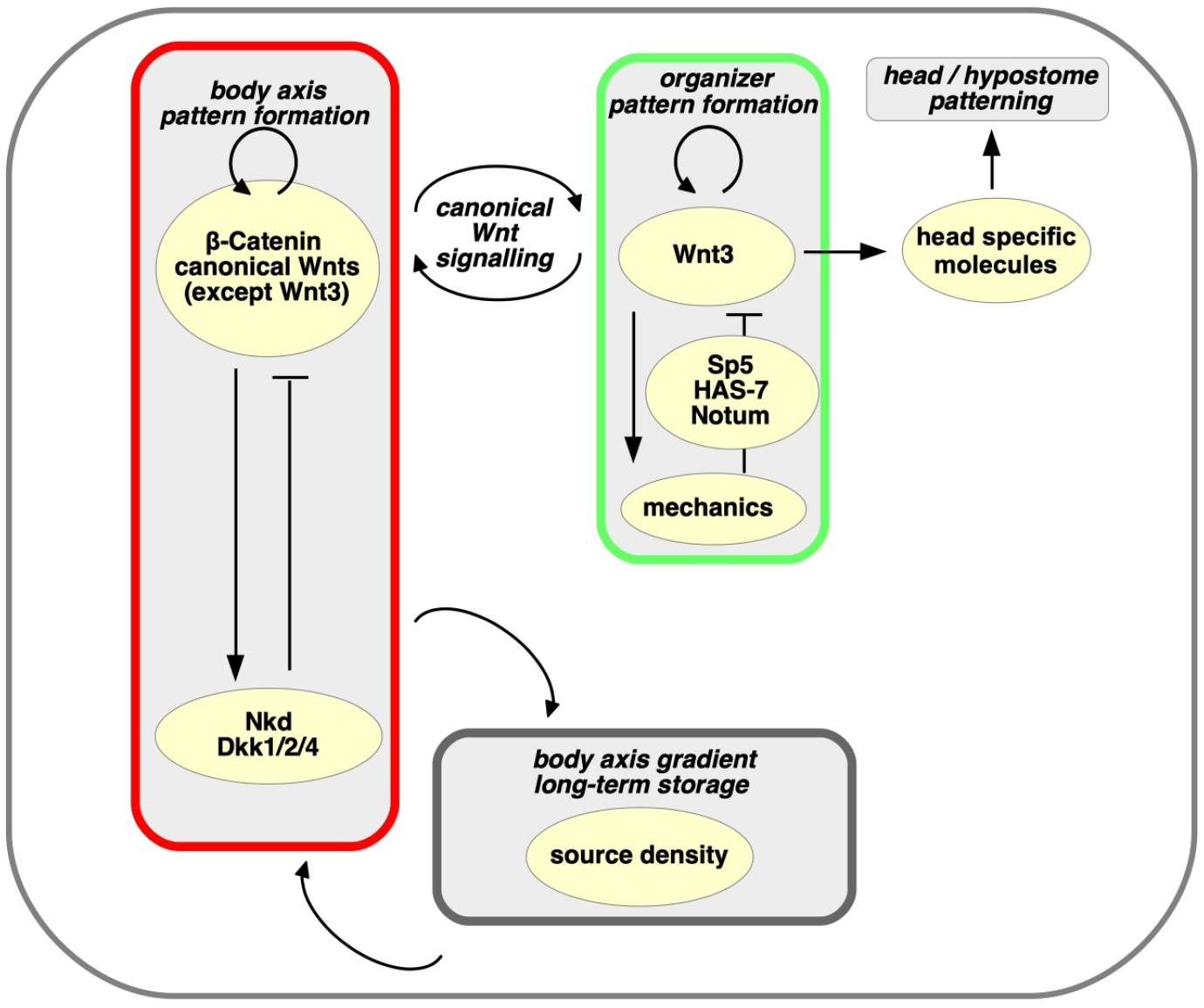
Possible molecular candidates for the double-loop model.

In summary, although there exist several known regulators of the Wnt3 and *β*-catenin dynamics, the related signaling systems are still not sufficiently understood to describe a complete working pattern formation mechanism without hypothetical components. Notably, all these mechanisms fit into the general concept of local activation and long-range inhibition (LALI). In this context, we replace the complex subsystems by simple pattern formation loops, using the inhibitor–activator model as a mathematical description of the LALI scheme. The consideration of patterning systems rather than individual molecules is supported by the fact that we obtained comparable results when replacing entire patterning units in our model by alternative mechanisms, i.e., replacing the activator-inhibitor loop by a mechanochemical one (see Supplementary Information S4). This is an important observation because we do not view our model as a tool to identify specific interactions at the molecular level, but rather as a tool to investigate the role of *β*-catenin versus Wnt3 in developmental and regenerative processes in *Hydra*.

In this study, based on a systematic analysis of previous and new experimental data, we proposed a revised two-scale mechanism and a corresponding mathematical model (double-loop model) for head and body-axis formation in *Hydra*. We conclude that there are two separate patterning systems for head (organizer) and body-axis formation, respectively, where Wnt3 is involved in a small-scale pattern formation system robustly forming the organizer, and *β*-catenin in interplay with other canonical Wnts leads to the *de novo* formation of the body axis. This paradigm shift has potentially important implications for the design and interpretation of previous and future experimental studies. This represents the first report of this novel concept, reflected by the fact that several recent papers investigating the molecular basis of *Hydra* patterning fail to distinguish between axis and head formation.

One of the main experimental implications of the proposed paradigm shift relates to the search for mechanisms leading to self-organized head/axis formation (such as molecules fitting the activator–inhibitor model): the search for a single pattern-forming mechanism is doomed to failure if there are actually two separate mechanisms, whose players have previously been mixed or equated, making interpretation of the experimental results difficult or impossible. The design and interpretation of future experiments should thus distinguish between the body axis *β*-catenin and the small-scale Wnt3 system. Indeed, candidate inhibitor mechanisms recently been identified for the Wnt3 patterning system (such as Sp5 (93), HAS-7 (103), and tissue stretch (98)). In contrast, details of the inhibitor molecules for the large-scale, *β*-catenin-driven body-axis system are still missing, although this patterning step is probably more fundamental and occurs before the formation of the head. Further studies are needed to investigate the roles of Nkd and Dkk1/2/4 molecules (2; 31; 58) in this respect.

Future experiments involving systematic knockdown or knockout of different genes could also be used to investigate the molecular candidates that interact with *β*-catenin and the molecules that are dispensable for self-organization of the body axis. Notably, recent experiments indicated that, in addition to diffusing chemicals, physical cues such as discrete cellular nearest neighbor interactions (11; 22; 40; 62; 75), tissue mechanics (96; 98), the extracellular matrix (91; 92), and bio-electrical processes (13) all actively contributed to pattern formation in *Hydra*. The integration of non-diffusive and diffusive signals thus offers a viable alternative to the classical Turing theory of *de novo* pattern formation and may have important consequences for experimental studies.

In summary, the molecular basis of canonical Wnt signaling established in previous decades (4; 35) clearly indicated a crucial role for Wnts in head and body-axis development, in both *Hydra* (34) and higher organisms (66). However, previous models failed to distinguish between body-axis and head formation in the context of canonical Wnt signaling. In contrast, employing a combination of mathematical modeling and simulations using previous and new experimental data, the current study provides the first evidence indicating that these two systems are controlled by two distinct self-organising processes. These two systems are coupled via the canonical Wnt signaling pathway, which ensures continuous alignment of the head with the body axis.

In 2012, based on over 40 years of research into *Hydra* pattern formation, Alfred Gierer noted that even a detailed knowledge of the molecules involved in *Hydra* pattern formation was not sufficient to explain the resulting spatial structures (25). We posit that the patterns that arise can be understood from an interdisciplinary perspective, integrating different chemical and physical processes and principles at the molecular, cellular, and tissue scales (12; 25). Further refinement of our models by integrating the results of different biological, biophysical, and biochemical disciplines will help to clarify the mechanisms driving self-organized pattern formation during development, as one of the key mysteries of biology.

## METHODS

### EXPERIMENTAL MODEL DETAILS

#### Hydra culture

Polyps were kept at 18 °C in *Hydra* medium (1 mM NaHCO_3_, 1 mM CaCl_2_, 0.1 mM KCl, 0.1 mM MgCl_2_, 1 mM Tris; pH 6.9) and were fed regularly three times a week with freshly hatched *Artemia salina nauplii*. The medium was changed daily. Animals were starved for 24 h prior to experiments. *Hydra vulgaris* AEP was used for all experiments, unless otherwise indicated. *wnt3*- overexpressing polyps carried a *Hydra actin* promoter driving the expression of *wnt3*. Transgenic *β-catenin* animals possessed additional copies of a *β-catenin* promoter-driven *β-catenin green fluorescent protein* fusion construct. Generation of both transgenic lines has been reported elsewhere (63). Phenotypical analysis and imaging was performed with bCat-Tg animals that carry an actin-driven bcatenin expression construct. The animals were a kind gift of Bert Hobmayer and have been initially described in Ref. (23).

#### RNA isolation and cDNA synthesis

Polyps were dissolved in 1 mL Trizol (Thermo Fisher) and 0.2 mL chloroform (Sigma-Aldrich). After centrifugation at 12,000 x g at 4 °C, the upper phase was transferred into a fresh tube and mixed with chloroform:isoamylalcohol at a ratio of 24:1. Samples were spun again as above and the upper phase was transferred into a fresh tube. RNA was precipitated with 0.8 volumes of pure isopropanol. After centrifugation, the pellet was washed again in 75 % v/v ethanol, dried in air, and taken up in nuclease-free water. RNA was digested with 1.5 U DNase I (Roche/Sigma-Aldrich) and subsequently inactivated according to the manufacturer’s instructions. The quality and concentration of the RNA were examined by 1 % w/v agarose (Invitrogen) gel electrophoresis in 1x TAE using a NanoDrop photometer. cDNA was transcribed using 1 *μ*g RNA and a sensiFAST cDNA Synthesis Kit (Bioline/Meridian) following the manufacturer’s instructions.

#### Whole-mount in situ hybridisation

ALP-treated animals were exposed to 5 *μ*M ALP in *Hydra* medium in the dark for 24 h. ALP was then removed by three washes in *Hydra* medium and the polyps were subsequently bisected at 50 % body length and allowed to regenerate for 72 h prior to fixation. Polyps for whole-mount in situ hybridisation were relaxed in 2 % w/v urethane in *Hydra* medium for 2 min at room temperature and subsequently fixed in 4 % w/v paraformaldehyde/ PBT at 4^°^C overnight. WISH was executed using DIG-labelled RNA probes (Roche), as described previously (8; 44).

#### Reaggregation assay

Wild type animals or polyps at 7-8 days post electroporation were amputated for head and feet at 80% and 20% body length, respectively. Body columns were rinsed three times in dissociation medium (3.6 mM KCl, 6 mM CaCl2, 1.2mM SO4, 6 mM Na-pyruvate, 4mM glucose, 12.5mM TES, 0.05 g/l rifampicin, 0.05 g/l kanamycin, 0.10 g/l streptomycin; pH 6.9) and subsequently incubated in dissociation medium for 5 minutes before tissue was gently dissociated by pipetting up and down with a glass pipette for 30 times. Remaining tissue was allowed to settle before the supernatant was transferred into a fresh tube. Procedure was repeated two additional times and the supernatant was spun down at 250 x g for 25 minutes at 10°C. Cell pellet was resuspended with fresh dissociation medium and partitioned to 0.4 ml tubes, which were once again spun down at 300 xg for 25 minutes at 10°C. Tubes were placed upside down in petri dishes filled with 75% dissociation medium/ 25% *Hydra* medium, in which the cell pellets were kept overnight at 18°C. On the next day cell pellets were gradually transferred to 100% *Hydra* medium. For inhibitor experiments, either 1 *μ*M iCRT14 (Sigma Aldrich /Merck) or 6.5 *μ*M LGK-974 (Selleckchem) were added when cell sorting was completed. As control, a similar volume of DMSO was used. Reaggregates were imaged at day 6 post dissociation using a Nikon SMZ-25 stereomicroscope.

#### siRNA-mediated knockdown

20 polyps were washed with water and subsequently placed in a 0.4 cm gapped cuvette (BioRad). Remaining water was removed and 200*μ*l siRNA mix was added to each cuvette. The final concentration of each mix was 3*μ*M, which was accordingly split up to 1.5 *μ*M when a combination of two siRNAs was used. When polyps were relaxed a single square wave pulse (BioRad Xcell pulser) at 240 V for 30 ms was applied. Chilled restoration buffer, consisting of 20% dissociation medium and 80% *Hydra* medium, was immediately added to electroporated polyps. Animals were then carefully transferred to petri dishes with chilled restoration buffer and allowed to recover for 2 days before the procedure was repeated for 2 more times. After the third pulse, animals were allowed to recover for 7-8 days with a daily feeding regime before they were used for reaggregation assays and validation by qRT-PCR. The sequences of the siRNA were purchased as duplexes from Merck using an UU-overhang. For simplicity, only the sense strands are given: siWntlessA 5′ GCCAAGACAAUUUCUUUAA 3′; siWntlessB 5′CCCUGCUACAGCCAAUAUA 3′; siWnt3A 5′ GGUGAUGCAACAUACGGAA 3′; siWnt3B 5′ AGAG-GCUAUAACGUUAAUA 3′; siGFP (control) 5′ GGUGAUGCAACAUACGGAA 3′. The success of the knockdowns was verified by means of qRT-PCR (Supporting Information S6).

#### quantitative RT-PCR

RNA was isolated as described previously (88). For cDNA synthesis, iScript cDNA Synthesis Kit (BioRad) was used with 1*μ*g of RNA according to the manufacturer’s instructions. cDNA was diluted 1:10 with nuclease free water and qRT PCR was performed with iTaq Universal SYBR Green Supermix (BioRad) according to the manufacturer’s instructions with a QuantStudio 5 Real-Time-PCR System (Thermo Fisher). For evaluation, all samples were normalized to EF1-a and fold change was calculated using the ΔΔCt method. The primer sequences were designed with the help of primer3 as exon-exon junction spanning primers.

Wnt1 fw 5′CCGATTATAGTCCAGAATCCGTTT 3′, rv 5′CACCAGCTGAAGTGATAGCGTAAAT 3′;

Wnt2 fw 5′CAAGTGGACTTATGAACAAACAAAA 3′, rv 5′AGAAACACAATGACAAATGCAAAT 3′;

Wnt3 fw 5′ATTACAACAGCCAGCAGAGAAAG 3′, rv 5′TTATCGCAACGACAGTGGAC 3′;

Wnt7 fw 5′ACCTGGTTGGCAATCACAAT 3′, rv 5′CTCGTTTTTACATAGATGACTGCTG 3′;

Wnt8 fw 5′AAGCGTTCTCCCACATTAATACTC 3′, rv 5′GCAACAATAGCATAAACATAAGCAG 3′;

Wnt9/10c fw 5′TGCTCTATAGTGTCTGGTTCTTTACT 3′, rv 5′TGAGCTAAACTTGCCGAGAATA 3′;

Wnt11 fw 5′CAAAGTGTAAACAAAGCTCCAGAAT 3′, rv 5′CTAGTGAAGAAGCTGCCAAAGAAT 3′;

Wntless fw 5′ATTGTTTAGCTGGCCAAGAA 3′, rv 5′TTTTGCTACAAATAACGAAATAGTGT 3′;

*β*-Cat fw 5′AAGTCAGCGTGCTAGAACAG 3′, rv 5′TGGTTCAGCAAGACGTTGAG 3′;

HAS-7 fw: GGATGTGAAATCAAATGGTTATGCT 3′, rv: TGATGAACTCATTCTTCGAAGATCG 3′;

Dkk124A fw 5′AGCCAAGTCTTGCAAAACGG 3′, rv 5′TCCCGGTTAAAGTTCCTCGT 3′;

Sp5 fw 5′ TCACCTCCAAGTCGTGTTCC 3′, rv 5′CAAGTAATGCCAACGGTGAATACT 3′;

ef1α fw 5′TATTGATAGACCTTTTCGACTTTGC 3′, rv 5′CTGTACAGAGCCACTTTCAACTTTT 3′

All samples were run in technical and biological triplicates.

#### Sphericity

The sphericity of the aggregates after 6 days of regeneration was determined from the 2D light microscopic images using the formula 4 * *pi* * *A*/*P*^2^, where *A* is the area of the aggregate volume and *P* is its perimeter. To approximate *A* and *P*, the images were loaded separately into Adobe Photoshop Elements Editor and the following steps were performed: (1) remove (blacken) all obvious tentacles and external loose cell structures; (2) possibly adjusting the contrast, subsequent contrast-based outlining the remaining aggregate (which stands out brightly against a black background), followed by manual correction of the outline if necessary; (3) count the corresponding pixels inside (Proxy for the area *A*) and the pixels of the border (for a border of 1 pixel wide – proxy for perimeter *P*).

#### Pairwise head distances

To measure and compare pairwise distances between heads, we analyzed light microscopic images of *N* = 10 budding polyps 3 days after Sp5-knockdown (taken from Supplementary Figure 6 of Ref. (93)) and from *N* = 15 *β*-catenin-overexpressing polyps. To ensure comparability, only polyps with three axes were analyzed. Distances were manually traced along the body axis in microscope images and normalized per polyp. As only 2D projections were analyzed, measurements approximate real 3D distances but may increase variance. Normality was tested using the Shapiro–Wilk test (*p* < 0.05 indicating deviation). Similar analyses were conducted based on the the double-loop model and for eight simulations per condition, starting from stochastic initial conditions. In simulated Sp5-knockdown, three Wnt3 spots were randomly selected per polyp for measurement. Since simulated body size was constant, no normalization was applied here. The underlying microscopic and simulated images can be found in Supporting Information S2.

### 1D/2D SINGLE-LOOP MEINHARDT MODEL

#### Model and its numerical simulation

The one-dimensional single-loop Meinhardt model is given by

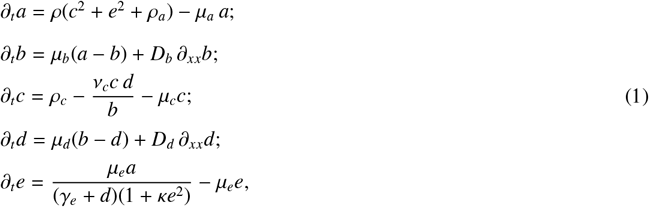

where *a* = *a*(*x, t*), *b* = *b*(*x, t*), *c* = *c*(*x, t*), *d* = *d*(*x, t*), and *e* = *e*(*x, t*) are unknown concentration functions (with *a* representing Wnt3), *ρ, ρ*_*a*_, *μ*_*a*_, *μ*_*b*_, *ρ*_*c*_, *ν*_*c*_, *μ*_*c*_, *μ*_*d*_, *μ*_*e*_, *γ*_*e*_, and κ are positive reaction parameters, *D*_*b*_ and *D*_*d*_ are diffusion coefficients, *x* ∈ (0, *L*) is the spatial variable, and *t* ≥ 0 is dimensionless time. More details of the model and the different components are given in Ref. (54; 55). The model is supplied with homogeneous Neumann boundary conditions:

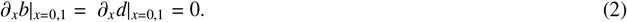

The model is discretized using the method of lines (MOL). We split the spatial interval (0, *L*) with the equidistant grid into *M* spatial nodes *x*_k_ = (*k* − 1) *h, k* = 1, …, *M*, where 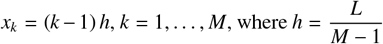. We approximate the Laplace operator with the three-point stencil:

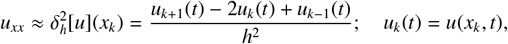

and arrive at the finite-dimensional system of 5*M* ordinary differential equations:

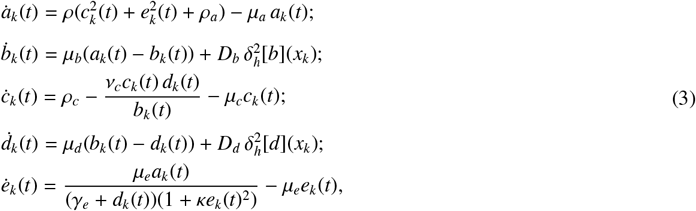

where *k* = 1, …, *M* and Neumann boundary conditions are taken into account by a central difference scheme (29):

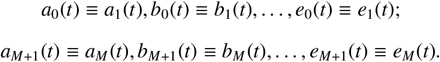

In our numerical experiments, we simulate the discretised system (3) using the small randomised perturbations of the spatially homogeneous steady state as initial data. All experiments are conducted using MATLAB, and numerical integration of the equations is performed using the explicit Dormand–Prince method (built-in ode45 routine). Additionally, we verify our results by integrating the system with the implicit TR-BDF2 (second order trapezoidal rule - backward differentiation formula) method (3).

### Heuristic algorithm for searching parameter values that belong to the Turing domain

To identify parameter values satisfying the conditions of Turing instability we develop a heuristic search algorithm. We consider system (1), group reaction parameters, and diffusion coefficients in the parameter vector ***θ*** and construct the objective function *f* (***θ***) using the following rule:

1. Consider system (1) without diffusion terms and find the spatially homogeneous steady state ***w***_0_(***θ***). This is done numerically using Newton’s method.
2. Set 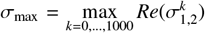, where 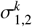 are eigenvalues of the linear operator *L*(***θ***), obtained by linearizing a spatially distributed system (1)-(2) around the steady state ***w***_0_(***θ***). Eigenvalues 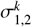 correspond to the respective eigenvalues of the Laplace operator −∂_*xx*_ with homogeneous Neumann boundary conditions (2), which are grouped in increasing order.
3. If 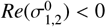 (system without the diffusion terms is linearly stable), then set *f* (***θ***) = σ_max_. Otherwise, reject the current value ***θ*** by defining *f* (***θ***) = −1*e*15.

Using the Differential Evolution algorithm (78), we perform a heuristic search in the parameter space to find a parameter vector ***θ***^*^ such that *f* (***θ***^*^) > 0.

### 2D/3D MATHEMATICAL MODELS AND SIMULATION DETAILS

#### Model geometry

The cell bilayer is at any time *t* approximated by a closed 2D surface ellipsoid Γ(*t*), embedded in 3D space. The evolution of Γ(*t*) is given by a diffeomorphic time-dependent representation 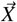, parameterized over the unit sphere *S* ^2^ ⊂ ℝ^3^. Thus, Γ(*t*) is the image of 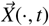 with 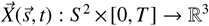 for a *T* ∈ ℝ_>0_. Local concentrations of different gene products at time *t* are given by a set of continuous functions Φ_*i*_, *i* = 1, …, *n*, on the deforming tissue surface Γ(*t*), defined as gene product concentrations per cell volume, Φ_*i*_(*t*) : Γ(*t*) → ℝ_≥0_. In order to achieve a consistent formulation with chemical processes being defined on *S*^2^ rather than on Γ, Φ_*i*_ are redefined accordingly. This can be achieved because we can identify material points 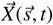 on Γ(*t*) with 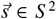, because 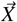 is smooth and bijective. Thus for each 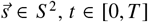, we define functions Φ_*i*_ : *S* ^2^ × [0, *T*] → ℝ_≥0_ by 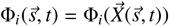.

#### Mathematical setting of the double-loop model

The model is given in terms of partial differential equations (PDEs) accounting for spatio-temporal interactions among the signaling factors defined on an evolving domain representing the tissue. Application of a continuous modeling approach is justified by the large number of cells (≥ 5 × 10^4^) in the system (82). The choice of model geometry is motivated by a realistic description of the tissue as a radially symmetrical ellipsoid undergoing small deformations due to changes in gene expression patterns. To describe its dynamics, we adopt a mechano-chemical modeling approach as proposed in Ref. (57; 59; 60). The signaling systems are described by reaction–diffusion equations describing defined on a curved 2D surface embedded in a 3D space. The geometry of the curved surface is assumed to evolve according to a gradient-flow of Helfrich-type energy, reflecting the fact that bending of the tissue away from a preferred local curvature is energetically unfavourable. Small tissue deformations are assumed, e.g., follow patterns of gene expression in the initial phase of tentacle development. In contrast to the fully coupled mechano-chemical model of (59), the main version of our presented model (for a mechano-chemical alternative, cf., below) does not consider any feedback between the mechanical properties of the tissue and the gene expression. In Ref. (59), we considered axis formation in mechanically oscillating *Hydra* re-aggregates of dissociated cells, which were described to require mechanics for patterning. In contrast, the current study focuses on axis vs. head formation in *Hydra*, which also comprises processes such as head regeneration, homeostasis, and budding. However, it remains unclear whether the mechanical stretching in aggregates is an artefact of the sphere-like reaggregates lacking a preformed mouth opening characteristic of intact animals, so the mechanical stress may not be essential for *Hydra* axis formation in general. Thus, for the sake of simplicity, we neglect the role of tissue stretching and consider a pattern formation process based solely on molecular interactions.

To demonstrate the robustness of the model results with respect to the choice of the pattern formation mechanism underlying each sub-system, we developed a second version of the model, relying on similar qualitative interactions between the different pattern formation systems but using different mechanisms for *β*-catenin and Wnt3 *de novo* pattern formation itself (cf., Supporting Information S3). Motivated by the increasing experimental evidence that tissue mechanics may actively influence axis formation (12; 45; 59), particularly *wnt3* expression (18) in *Hydra*, we replaced the activator–inhibitor scheme by a a mechanochemical model where *wnt3* expression and tissue curvature are coupled in a positive feedback loop (57). Furthermore, for the *de novo* formation of the body axis, we replaced the Turing mechanism by the mutual inhibition model recently presented by (58) (further details are given in Supporting Information S3).

#### Interactions between molecule groups

In the double-loop model, we considered eight different groups of molecules Φ_*i*_ (denoted in the following, if known, by its most prominent representatives), where five groups build the core pattern formation system explaining axis vs. organizer formation (namely *β*_*cat, β*_*cat*_*ant*_, *Wnt*3, *Wnt*3_*ant*_, and *S D*) and three components are augmented to include further head and tentacle development (namely *Head, S D, Tent*, and *Tent*_*ant*_). However, the latter were not required for all the main statements of this work, given that the interplay with the tentacle system was not the main focus of the study. An overview of possible molecular candidates is given in Table 1. Here, *β*_*cat* and *β*_*cat*_*ant*_ represent the activator and inhibitor associated with the body-axis patterning system involving nuclear *β*-catenin. *Wnt*3 and *Wnt*3_*ant*_ are the activator and inhibitor of the Wnt3 organizer pattern formation system, respectively. *Head* is one of several head-specific gene products downstream of the organizer (such as one of the multiple head-specific Wnts (67)), *SD* denotes the source density, and *Tent* and *Tent*_*ant*_ describe the activator and inhibitor of the tentacle system, respectively. The PDE system describing interactions among these components is given in Eqs. (4)-(8) (core system) as well as Eqs. (9)-(11) (head and tentacle system), where Δ^Γ^(.) denotes the surface Laplace (Laplace–Beltrami) operator, and *d*_*t*_ stands for time-derivative. Specifically, Eqs. (1)–(2) describe the body-axis pattern formation system, Eqs. (3)–(4) describe the organizer pattern formation system, Eq. (5) is for the source density, Eq. (6) describes the equation for *Head*, and the tentacle system is given by Eqs. (7)–(8).

**Table 1.** Model variables and possible molecular representatives.

| Variable name | Explanation |
| --- | --- |
| $\beta\_cat$ | Nuclear β-catenin, possibly linked to other factors such as Tcf and/or other Wnts than Wnt3 (e.g., Wnt9/10c) (44; 88) |
| $\beta\_cat_{ant}$ | $\beta\_cat$ antagonist (presumably linked to Dickkopf1/2/4-C (2; 54; 58) and/or HyNkd) |
| $Wnt3$ | Wnt3 |
| $Wnt3_{ant}$ | Wnt3 antagonist (related to Sp5 (93), HmTSP (46), or Notum signaling) |
| $Head$ | Head-related factors downstream of Wnt3 (e.g., multiple Wnts (44)) |
| $SD$ | Source density (long-term storage of head-forming potential (88)) |
| $Tent$ | Tentacle activator (may involve HyAlx (74), Wnt8, and Wnt5 (67) or BMB5-8b (68)) |
| $Tent_{ant}$ | Tentacle activator antagonist (unknown) |

Although *β*-catenin is a non-diffusive molecule, we modeled the pattern formation system of *β*-catenin using a system of reaction–diffusion equations, because it might interact with other (possibly diffusing) ligands, such as Wnt9/10c. The diffusion-based model can also be seen as an abstract representation of a more complex pattern formation model in which effective short-range cell-to-cell communication may be realized through interactions with Wnt signaling (4). A mechanism of pattern formation exhibited by components of intracellular signaling coupled to a single diffusive component has been demonstrated in a series of mathematical works (32; 51). The desired pattern formation mechanism can be also realized by the interplay between *β*-catenin and mechanical cues. The formation of morphogen patterns in the absence of lateral chemical diffusion has recently been demonstrated in simulation studies based on a mechano-chemical model (14). Considering several possible molecular mechanisms that could explain the formation of the *β*-catenin patterns, we formulated the current model in terms of a reduced mathematical metaphor for such mechanisms. A similar modeling strategy was applied in the seminal works of Meinhardt (53; 54). In particular, we assumed that *β-catenin* expression requires the source density (54; 88) and, due to canonical Wnt signaling, is additionally supported by Wnt3.

The extension of the current model compared with previous models includes a separate activator–inhibitor system for the Wnt3 organizer system and the additional variable *Head*, which represents the head-specific molecules expressed downstream of the organzser, in order to fine-scale the pattern of the *Hydra* head region (e.g., multiple Wnts (44)). The activator–inhibitor system for Wnt3 is modeled as a small-scale pattern formation system with saturation of its own expression, motivated by the sizes and spacing of the (transient) Wnt3 spots after ALP treatment. The production of Wnt3 is assumed to require *β*-catenin. Wnt3 again activates head-specific molecules (*Head*), but only above a certain threshold, *thresh* = 5.0. We introduced this threshold to account for the assumption that head construction only starts if the organizer is well established. *Head* is again assumed to negatively influence the tentacle system. This negative influence of the *Hydra* hypostomal region on the tentacle system has been demonstrated in different experiments (68; 74; 81).

The large total number of variables/fields in the presented model may lead to the criticism that the model includes “…so many parameters that they can made to fit anything…” (47). Notably however, only equations (1)–(5) are required to reproduce the main results of this study. In addition, most parameters in Eqs. (1)–(8) are not critical for the qualitative simulation results presented in this study; most of them control specific properties of one of the three considered *de novo* pattern formation systems, such as spatial scaling in general, spacing between appearing maxima, or the condition required to start the *de novo* patterning process. Most of these parameter have been adopted from Ref. (53; 54). They are not within the focus of the current study and do not critically influence our main results, given that general interactions between the different pattern formation systems in *Hydra* are considered, rather than the details or the *de novo* pattern formation systems themselves (cf., Supporting Information S4).

The detailed molecular mechanisms responsible for the latter are still elusive and need further research, but are beyond the scope of the current study. The activator–inhibitor model has thus been used just as an appropriate ‘toy-model’. The main focus of this study was therefore the aspects of *Hydra* patterning that do not depend on the specific choice of *de novo* patterning mechanisms, including the existence and interplay of the different patterning systems, rather than the identification of specific molecular interactions that could fit to a specific pattern formation mechanism.

We therefore believe that our main conclusions were largely unaffected by most of the model parameters. Because the simulation times required for a systematically numerical robustness analysis for our models are not feasible, we demonstrated this robustness by developing and simulating an alternative model relying on similar proposed scales and interactions between the distinct pattern formation systems, but realizing *de novo* patterning for each single system in completely different ways. More details are given in the Results and Discussion and in Supporting Information S4.

Core pattern formation system:

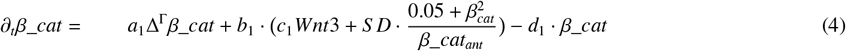

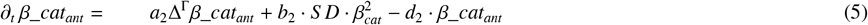

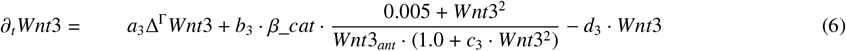

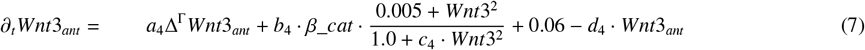

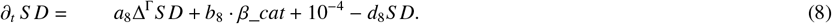

Additional equations for further head and tentacle formation:

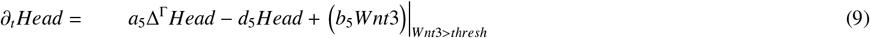

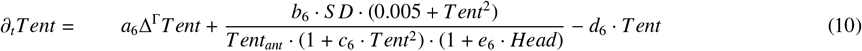

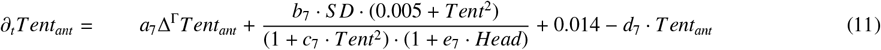

#### Single-loop model

The single-loop model considered is based on the idea proposed by Hans Meinhardt to model Wnt3 and *β*-catenin patterns in an extended single-loop activator-inhibitor system with a threshold mechanism linking localized patterns of Wnt3 to broadly supported patterns of nuclear *β*-catenin (54; 55). The original idea was implemented by means of a system of reaction-diffusion- ODE equations (cf., Supporting Material S3). Simulations of this model show emergence of irregular spikes, similar to those previously observed in a class of reaction-diffusion-ODE systems (32; 50; 70), see Supporting Information S3 for more details. Emergence of such singular patterns is due to the lack of diffusivity in some model components, what may be linked with the observed concentration effects (emergence of unbounded spike solutions (70)). This observation motivated us to replace system (1)–(8) by some prototype models of the core Mainhardt’s idea of the single-loop model. To this end, we replaced Eqs (2)–(3) by one equation for Wnt3 (cf., Eq (9) below) assuming *wnt3* expression above a certain critical threshold of *β*_*cat*. All other equations, parameters and numerical routines are identical to the double-loop model. Different scenarios of a single-loop model are presented in Figure. 1 left panel.

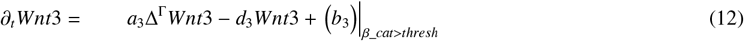

#### Tissue curvature

The chemical/molecular equations Eqs. (1)–(8) are augmented by a set of equations representing the deforming tissue surface. Notably, we treat the tissue as purely elastic, and elastic tissue deformations were based on minimization of the Helfrich free energy (33), given by

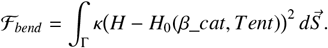

Here, *H* is the mean curvature, κ the bending rigidity, and *H*_0_ the spontaneous curvature (57; 60). *H*_0_ represents the locally preferred tissue curvature, which again may depend on local morphogen concentrations. Here, we assume *H*_0_(*β*_*cat*_*ant*_, *Tent*) = 0.5 · *β*_*cat*_*ant*_ + 15. · *Tent*, based on the observation that local tissue evaginations can be observed during both budding and tentacle formation (1; 67). Local area-conserving evolution of the deforming *Hydra* tissue is finally given by the *L*^2^-gradient flow of the total energy. Further details are given in Ref. (60).

The model proposed in this study focuses on the interplay of pattern formation feedback loops in the biochemical signaling system, and simulated changes in tissue curvature are thus only a read-out of biochemical morphogen concentrations. We include the resulting evolution of the tissue geometry in our model (through a respective PDE describing tissue mechanics), because even the dynamics of a purely biochemical model depend on the domain size and geometry that may influence the appearance and spacing of the organising centers. Various theoretical studies have demonstrated that Turing-type patterns depend directly on the underlying geometry leading, e.g., to spots on the body and stripes on the tail during animal coat patterning. We therefore consider that the interaction of biochemical patterning and tissue geometry, which approximates the real geometry, is important for obtaining realistic patterns. This allows for a more realistic description of tentacle formation in our model, which can again be used for further model validation (e.g., simulating ALP treatment). However, exemplary simulations without tissue deformations reveal only minor differences, suggesting that tissue geometry plays only a subordinate role (Supporting Information S3).

We previously demonstrated (44; 67) that Wnt5 and Wnt8 acted downstream of canonical Wnt 3 and Wnt9/10 signaling. Wnt5 acts as a non-canonical Wnt signal during tentacle formation, and non-canonical Wnts are known to be involved in planar cell polarity, which is in turn closely linked to changes in cell geometry. Downstream of non-canonical Wnt signaling during tentacle outgrowth, *Hydra* Alx-spots change from spots to rings (74), probably caused by the interplay between chemistry and geometry. Nevertheless, although there is likely to be an interplay between chemical signaling and the underlying evolving tissue geometry, strong geometric changes were not considered in the present study because they seemed to be irrelevant to the mechanism of axis formation per se.

#### Simulation method

For simulations of the model, we use the finite element library Gascoigne (6), approximating the fourth order PDEs in a mixed formulation. For spatial discretization, we use linear finite elements (including local mesh refinement in areas with strong local curvatures), and for time discretization, we use a semi-implicit Euler scheme (including an adaptive time-step scheme). Further details of the computation scheme are provided in (57; 60).

#### Parameters and initial conditions

For simulations of the undisturbed system, most of the parameters are adapted from Ref. (53; 54). For the unperturbed system, we use: *a*_1_ = 9 × 10^−5^, *b*_1_ = *d*_1_ = 3 × 10^−3^, *a*_2_ = 1.1 × 10^−2^, *b*_2_ = 3 × 10^−3^, *d*_2_ = 4 × 10^−3^, *a*_3_ = 2.25 × 10^−5^, *b*_3_ = 4.8 × 10^−2^, *c*_3_ = 1.8 × 10^−3^, *d*_3_ = 1.2 × 10^−2^, *a*_4_ = 0.9 × 10^−3^, *b*_4_ = 0.72 × 10^−1^, *c*_4_ = 1.8 × 10^−3^, *d*_4_ = 1.8 × 10^−2^, *a*_5_ = 5 × 10^−4^, *d*_5_ = 1 × 10^−2^, *b*_5_ = 1, *a*_6_ = 7.5 × 10^−5^, *b*_6_ = 2 × 10^−2^, *c*_6_ = *e*_6_ = 1 × 10^−1^, *d*_6_ = 2 × 10^−2^, *a*_7_ = 3 × 10^−3^, *b*_7_ = 3 × 10^−2^, *c*_7_ = *e*_7_ = 1 × 10^−1^, *d*_7_ = 3 × 10^−2^, *a*_8_ = 1.1 × 10^−4^, *b*_8_ = *d*_8_ = 1 × 10^−4^. To approximate the *Hydra* shape, initial conditions for *X*_1_, *X*_2_, and *X*_3_ are parameterized over the closed 2D unit-sphere *S* ^2^ embedded in 3D space with *X*_1_(*t* = 0) ≡ *X*_2_(*t* = 0) ≡ 0 and *X*_3_(*t* = 0) = 4 · *s*_3_, thus leading to stretch in the direction of *s*_3_ (given that *s*_1_, *s*_2_, *s*_3_ are Eulerian coordinates of the *S* ^2^ surface). For chemicals, we use a stochastic initial distribution based on the standard random generator provided by C++. The only exception is the source density, which is provided with an initial gradient, namely via *S D*(*t* = 0) = 4.0 · (*exp*(*s*_3_)/*exp*(1)). Thus, in all simulations, chemical patterns follow stochastic initial conditions, with only the geometric and chemical body-axis gradient initially provided. For simulations including ALP treatment, the initial conditions for the source density are changed by adding an offset by *S D*(*t* = 0) = 2.0 + 4.0 · (*exp*(*s*_3_)/*exp*(1)). *sp5* knockdown is simulated by increasing degradation of *Wnt*3_*ant* by multiplying *d*_4_ by the factor 1.7, and overexpression of *β-catenin* and *wnt3* are realized by adding the constant *c* = 0.003 to the right-hand side of the corresponding equation.

## DATA AND CODE AVAILABILITY

This study did not generate any of the dataset types for which deposition in a public repository is mandated by the journal. Therefore, no data repository is required. All relevant data supporting the findings of this study are available within the manuscript and Supplementary Information. The FEM basis code for the 2D/3D simulations is available at *https://www.uni-kiel.de/gascoigne/* (cf., (6)); specific codes for Hydra simulations are available from the lead contact upon request (Moritz Mercker); corresponding parameters are given in the Methods section.

## ACKNOWLEDGEMENTS

This work was supported by Deutsche Forschungsgemeinschaft (DFG) under Germany’s Excellence Strategy EXC-2181/1 - 390900948 (the Heidelberg STRUCTURES Excellence Cluster) and SFB1324 (B05 to AM-C and MM, B07 to SÖ, and A05 to TWH and AT). We thank Yukio Nakamura for preparation of the actin::Wnt3 pBSSA-AR vector and initial transgenic *Hydra vulgaris* AEP strains (DFG-FOR 942; TWH).

## AUTHOR CONTRIBUTIONS

MM designed the conceptual models and integrated them into the computational 3D framework, performed 3D simulations, carried out the systematic experimental review, and developed the main research questions. AM-C and MM designed, guided and integrated the supporting experimental, numerical and analytical work. MM and AM-C wrote the manuscript with input from all authors. FV performed the analytical studies and AK performed the numerical 1D analyses. TWH and SÖ conceptually accompanied the experimental work and interpretation. AT carried out the main experimental work, supported by SH and TL. AM-C, TWH, and SÖ conceived the experimental-theoretical interplay and were in charge of overall direction and planning.

## SUPPLEMENTAL MATERIALS

- Pdf-document with text and figures (SI.pdf)

