## Supporting Material for "Two interconnected patterning loops are required for body axis and head organizer formation in *Hydra*"

### Contents

**1 S1: Details of previous experiments indicating distinct intrinsic and pattern formation functions of $\beta$-catenin and Wnt3 — 2**
- 1.1 Regeneration and grafting experiments — 2
- 1.2 Developmental viewpoint — 2
- 1.3 Single-gene expression studies — 3
- 1.4 Transcriptomic analyses and signaling pathways — 3
- 1.5 Molecular manipulations and multi-headed phenotypes — 3
- 1.6 Mutual positive influence — 4

**2 S2: Pairwise head distances in manipulated experimental and simulated *Hydra* polyps — 4**

**3 S3: Numerical pattern analysis for the single-loop Meinhardt model — 5**

**4 S4: Variations of the double-loop model — 6**

**5 S5: Expression patterns of *Naked cuticle (Nkd)* — 7**

**6 S6: Knockdown validation — 8**

### 1 S1: Details of previous experiments indicating distinct intrinsic and pattern formation functions of $\beta$-catenin and Wnt3

In this chapter, we summarize in more detail previous experimental studies that deal with the differences between Wnt3 and $\beta$-catenin. In doing so, we do not explicitly distinguish (unlike in the main manuscript) between the intrinsic and pattern formation function of these molecules, since this distinction is not strictly possible in most of the listed studies.

#### 1.1 Regeneration and grafting experiments

Several organism-based studies have indicated that head and body-axis formation in *Hydra* are established (or at least realized) by two different systems / mechanisms acting at two different spatial scales. Although both *Hydra* head and body-column tissue can lead to ectopic head and axis formation when grafted into a host polyp (5; 10), only the apical part of the head (the hypostome) exhibited organizer capacity, transmitting a signal that induced the host tissue to form a secondary axis (10). Grafted body-column tissue did not stimulate the surrounding tissue to a similar extent, but self-organized into a secondary axis composed mainly of the transplanted tissue (5; 10; 12; 13). Furthermore, tissue from the apical tip induced a secondary axis more frequently(29) than grafts from other parts of the body (e.g., (34; 35)). Moreover, even pieces smaller than the hypostome, i.e., clusters of approximately 10 epithelial cells derived from the regenerating hypostome, induced head-organizing centers in reaggregates (53). All these observations suggest that the processes of axis and head formation originating from the apical tip (small-scale structure) vs. body-column tissue (large-scale structure) must involve different signaling factors (10; 35).

#### 1.2 Developmental viewpoint

The existence of two patterning systems acting at different scales to control robust formation of the *Hydra* body plan is also supported from a developmental perspective. First, *de novo* pattern formation at the scale of the body axis is indicated by the self-organization process during regeneration in *Hydra* aggregates. The resulting body-axis gradient is in accord with the experimental observation that transplantation of tissue pieces from the head region is more likely to induce a secondary axis than grafts from more basal regions. The presence of this 'head-activation gradient' was confirmed by several experiments (35). This property has also been referred to as 'head inducing potency' (19), gradient of 'head formation capacity' (10), ' -competence' (53), '-potential' (24), '-positional value' (1; 61), and 'source density' (18; 38).

In addition, this head-activation gradient has to be subsequently translated into fine-scale patterning in the *Hydra* head region. Regeneration experiments indicated that the location of head vs. foot formation depended on relative, rather than absolute levels of the head-activation gradient (3; 4; 43). Hence, 'direct' translation of the gradient into fine-scale patterns, e.g., based on a threshold mechanism as originally proposed by Lewis Wolpert (60; 61), is relatively unlikely. In addition, such a mechanism would require the detection of very small concentration differences between adjacent cells (37).

In contrast, a partially independent small-scale pattern formation system for head patterning, activated by high levels of components of the large-sale system, would provide a much more robust strategy to realize the transition from large- to small-scale patterns without critical/sensitive threshold dependencies. This would ensure correct localization of head patterns, without depending on absolute levels of components in the large-scale system.

At a molecular level, the large-scale head-activation gradient was shown to co-localize with nuclear *$\beta$-catenin/T-cell factor* (*Tcf*) expression, which appears to induce the 'head organizer', the latter defined by *Wnt3* expression (5; 6; 11; 25) and always localized to maximal (or sufficiently high) levels of nuclear $\beta$-catenin (25; 53). The organizer spot is finally assumed to control the stepwise construction of the *Hydra* head, including the tentacles (3; 6; 8; 10; 31).

In summary, instead of the potentially non-robust direct transformation of the large-scale gradient into small-scale patterns, the large-scale body-axis gradient seems to be involved in mutual positive feedback with another, small-scale patterning system defining the head organizer. Robustness is thus achieved via redundant mechanisms acting at different scales, representing a well-known principle in various biological contexts (57).

#### 1.3 Single-gene expression studies

*In situ* hybridization- and green fluorescent protein-based reporter gene studies revealed that, although *β-catenin* and *Wnt3* were co-localized with respect to their maxima, they were expressed at different spatial scales: *β-catenin* (and accordingly *Tcf*) showed more-diffuse, longer-ranging expression patterns, whereas expression of *Wnt3* and all other *Hydra Wnt* genes always occurred in small spots (11; 17; 25; 28; 31; 52; 53). The activities of these two factors differed not only in terms of their spatial patterns, but also in their dynamics. With the exception of injury-induced *Wnt3* expression (56), experiments on intact polyps suggested that *Wnt3* was expressed at later stages than *β-catenin. Wnt3* expression was distinctly delayed compared with *β-catenin* during embryogenesis (16) and budding; whereas *Wnt3* expression commenced with the appearance of the first bud protrusions (25), *β-catenin* expression was already evident prior to the first visible deformations (28; 58). Indeed, inhibition of *β-catenin* prevented bud formation, including the expression of several bud- and organizer-specific genes (58).

In summary, the results of these spatial and temporal expression studies suggest that the activities of both molecules are controlled by different mechanisms acting at different spatio-temporal scales, with *Tcf/β-catenin* frequently preceding *Wnt3* expression.

#### 1.4 Transcriptomic analyses and signaling pathways

Recent progress in our understanding of canonical Wnt signaling in different animals provides a direct link among secreted Wnt molecules, nuclear β-catenin, and Tcf transcription factors (2; 30; 45). This has led to the widespread assumption that β-catenin and Wnt3 must act within the same pattern formation system, with secreted Wnts at the core of this system (58). However, the observation that *Wnt3* expression requires nuclear β-catenin does not imply that the body-scale pattern formation system coordinated by β-catenin relies on Wnt3. Indeed, while Wnt3 function is associated with the head organizer (5; 6; 11; 25), β-catenin-dependent signaling is involved in diverse molecular, non-molecular, intrinsic, and extrinsic processes. Its role is thus versatile and fundamental: it is required for the early regeneration response and pre-patterning (22), and controls the expression of approximately 1000 different genes, including several components of the bone morphogenetic protein, Notch, and transforming growth factor-β pathways, several transcription factors, as well as genes influencing cell-cell adhesion, cytoskeletal components, and the extracellular matrix (46; 58). Regarding *Hydra* axis formation, the Nodal–Pitx cassette was demonstrated to act under the control of β-catenin (58). Moreover, β-catenin appears to be a required co-factor for establishing bi-radial budding asymmetry (58). In addition to its role as an important transcriptional co-factor in cell nuclei, the membrane-associated distribution of β-catenin in other body regions indicates a function in *Hydra* tissue mechanics (14; 25; 26; 28), such as cell-cell adhesion (44). Furthermore, *β-catenin* expression levels have recently been linked to extracellular matrix elasticity (54), which in turn plays important biochemical and mechanical roles during pattern formation and regeneration.

External biotic and abiotic environmental cues have also recently been shown to affect β-catenin-mediated signaling and pattern formation (7; 51). The complexity of β-catenin-related processes suggests that β-catenin-dependent *Wnt3* expression is only one of several pattern formation loops activated by gene expression downstream of β-catenin (45; 58), thus possibly making *Wnt3* expression dispensable in terms of body-axis formation, where the latter might instead with other molecules (such as Dkk molecules or other Wnts). This dispensability is supported by recent experiments in which *Wnt3* expression (and thus head formation) was suppressed but foot development occurred, possibly indicating preceding self-organization of a body axis in *Hydra* aggregates (15). In turn, recent studies suggest that Wnt3 patterning does not strictly rely on β-catenin signaling: *Wnt3* expression was not inhibited and a normal hypostome was regenerated when β-catenin was pharmacologically inhibited via iCRT14 (even if these animals did not regenerate complete/functional heads) (50).

#### 1.5 Molecular manipulations and multi-headed phenotypes

Recent advances in molecular techniques have opened the way for the analysis of *Hydra* phenotypes resulting from body-wide overexpression in transgenic animals or knockdown of single genes using RNA interference (33; 59). Notably, *β-catenin* knockdown (55) and overexpression (17) have been studied in *Hydra*. *Wnt3*-overexpressing transgenic polyps are also available (62), and knockdown of putative Wnt3 antagonists has recently been carried out. Antagonists can either interfere on the level of transcriptional regulation or by inhibiting the formation of receptor–ligand complexes, such as the Wnt antagonist Dickkopf (Dkk 1/2/4) (1; 21; 41), Sp5 (55), and the Wnt-specific protease HAS-7 (62). As expected, overexpression (direct or indirect resulting from inhibition of an inhibitor) of either *β-catenin* or *Wnt3* produced phenotypes with multiple heads. However, a more detailed examination of these phenotypes allowed the distinction of two different categories. (See main manuscript for more details).

##### 1.6 Mutual positive influence

Despite the many above-mentioned studies that point to differences in the intrinsic and pattern formation function, there is much evidence that both molecules (in line with our knowledge of canonical Wnt signaling) are linked in a positive loop. In in particular, both expression levels were highest at the oral end of the *Hydra* body column and at the tips of developing buds (4; 5; 10; 17; 25; 27; 31). Furthermore, pharmaceutical or transgenic induction of nuclear $\beta$-catenin induced transient increases in the number of HyWnt3 spots (11; 17; 20; 21). Several studies also showed that HyWnt3 mRNA expression is induced by (or dependent on) $\beta$-catenin (45; 46; 55). On the other hand, it has been shown that the organizer (characterized by local *Wnt3* expression) induces the formation of a new body axis (10) (which in turn is strongly associated with a gradient of nuclear *beta*-catenin (17; 25)).

#### 2 S2: Pairwise head distances in manipulated experimental and simulated *Hydra* polyps

Experimental measurements underlying Fig. 2 (c)–(f) in the main manuscript are shown in1.

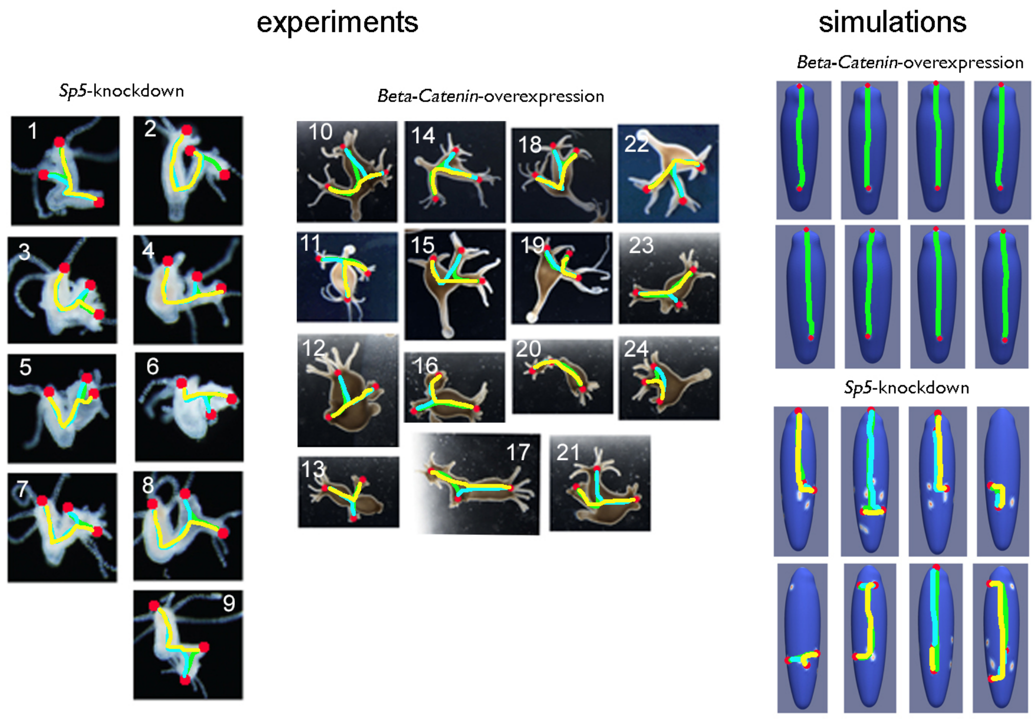


**Supplementary Figure 1.** Left-hand side: Measurements of pairwise distances (colored lines) between heads (red dots) based on *Sp5*-knockdown (left) vs. *β-catenin*-overexpressing (middle) polyps. Polyps on the left-hand side taken from Supplementary Figure 6 (3 days after siRNA treatment) of Ref. (55). Right-hand side: simulation results for *Sp5*-knockdown (below) vs. *β-catenin* overexpression (above) based on the double-loop model.

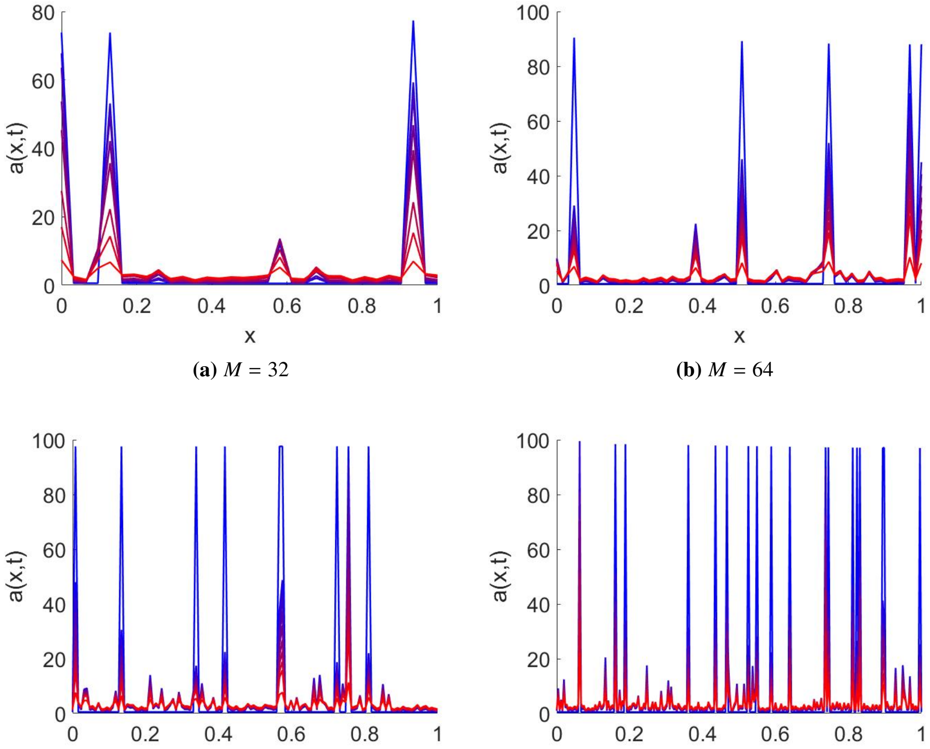


**(a)** $M = 32$ **(b)** $M = 64$ **(c)** $M = 128$ **(d)** $M = 256$

**Supplementary Figure 2.** Numerical solution of MOL system showing simulated Wnt3 concentrations for different values of grid resolution $M$. The transition from red to blue corresponds to increasing time.

### 3 S3: Numerical pattern analysis for the single-loop Meinhardt model

In the following, we numerically investigate the single-loop model proposed by Hans Meinhardt to explain patterns of Wnt3 and $\beta$-catenin in *Hydra* (38; 39). We demonstrate that the system does not describe robust patterns. The spatially heterogeneous solutions emerging due to Turing-type instability have the form of irregular infinitely growing narrow spikes, similar as observed previous in a class of reaction-diffusion-ODE models (23; 36; 47).

We begin from identifying parameter values satisfying the conditions of Turing instability. Using our heuristic algorithm, we find a parameter vector $\boldsymbol{\theta}^*$ such that $f(\boldsymbol{\theta}^*) > 0$, which means that the point belong to the Turing domain. We then simulate the MOL system for increasing values of the grid resolution $M$. As initial data, we consider small random perturbations of the homogeneous steady state $\boldsymbol{w}_0(\boldsymbol{\theta})$. All the simulations show convergence of the solution to irregular narrow spikes. The number and amplitude of spikes grows with increasing grid resolution $M$ (see Supplementary Figure 2).

### 4 S4: Variations of the double-loop model

In order to demonstrate that the presented results do not depend on the particular choice of the inhibitory mechanism or model parameters, we consider an alternative version of the double-loop model ('alternative model 1') using similar qualitative interactions between the different pattern formation systems, but replacing the Turing-type activator–inhibitor sub-models for $\beta$-catenin and Wnt3 by alternative mechanisms. Notably, for the *de novo* axis formation process, we assume the 'mutual inhibition' mechanism describing the interplay between two different HyDkk molecules and the canonical Wnt signaling complex. This mechanism has recently been demonstrated to not only recapitulate *de novo* axis formation, regeneration, and grafting experiments, but also explained the interplay between wound response and pattern formation initiation in *Hydra* (41). With respect to the small-scale Wnt3 patterning mechanism, we replaced the activator–inhibitor description by the mechano-chemical mechanism (40; 42), based on the increasing evidence indicating that mechanical processes may play an active role during patterning in *Hydra* (9; 15; 32; 42; 48; 49).

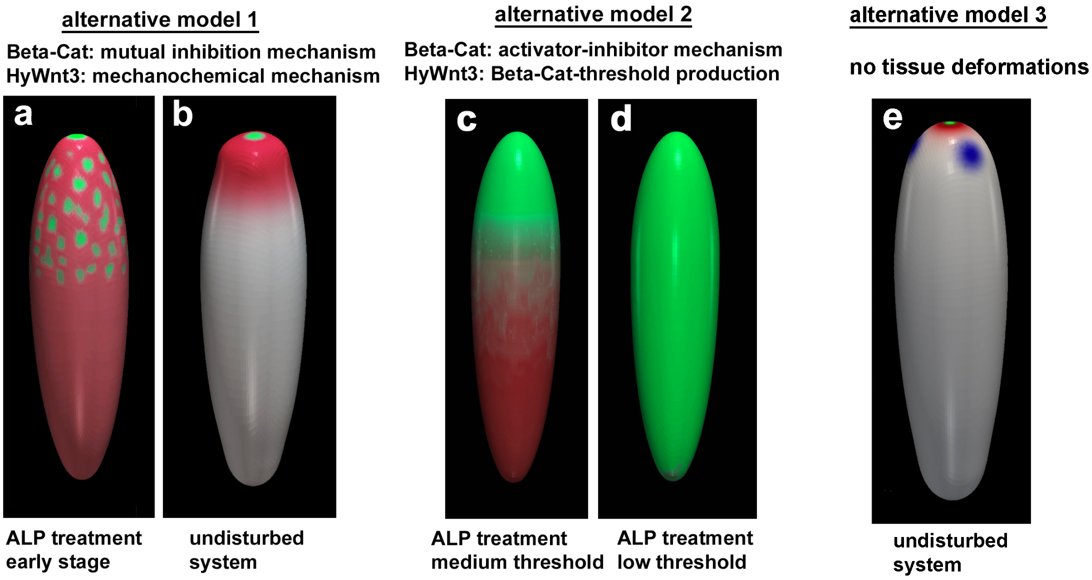


**Supplementary Figure 3.** Simulations of *Wnt3* expression (green) and $\beta$-catenin expression (red) for three alternative models and different treatments compared with the undisturbed scenario. In 'alternative model 3', blue corresponds to the tentacle activator, which is not shown in the other simulation snapshots.

The simulation results for alternative model 1 for the early stages of ALP treatment and the undisturbed system are shown in Supplementary Figure 3 (left-hand side). This model also recapitulates key experimental results indicating the existence of two separate pattern formation systems acting at different scales, namely the transient expression of multiple *Wnt3* spots following ALP treatment. To further demonstrate that this behavior cannot be observed in the context of 'threshold models', i.e., the idea that observed scale differences are induced by a mechanism where *Wnt3* is produced above a certain threshold of $\beta$-catenin, we numerically investigate a second type of model ('alternative model 2'). This model is similar to the double-loop model presented in our study, but instead of a self-contained patterning system for Wnt3, *Wnt3* expression is model by a threshold function of $\beta$-catenin. The simulation results for alternative model 2 for the early stages of ALP treatment show either no (c.f., main manuscript) or broad (Supplementary Figure 3 c–d) expression patterns of Wnt3, depending on the threshold values. This behavior strongly contradicts the experimental observations on the formation of multiple transient *Wnt3* expression spots (17).

### 5 S5: Expression patterns of *Naked cuticle (Nkd)*

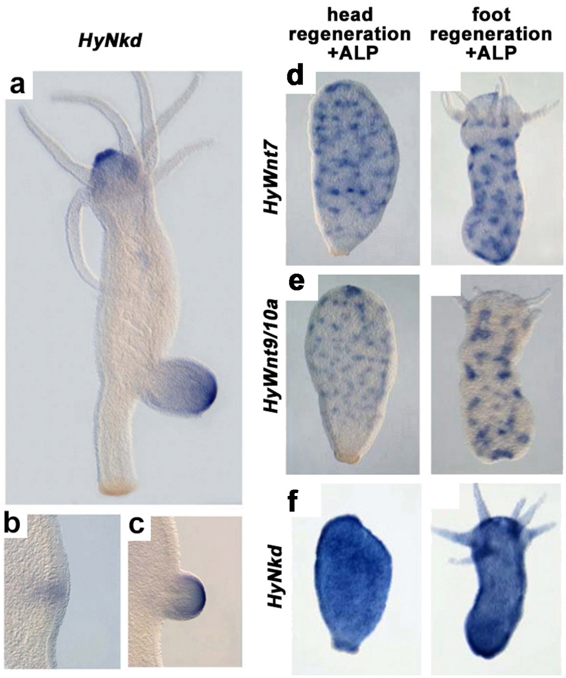


**Supplementary Figure 4.** Experimental comparison of large-scale diffusive expression patterns and small-scale spotty expression patterns of different genes suggesting their involvement in the two different patterning systems. (a–c) Analysis of large-scale expression patterns of HyNkd in adult polyps, suggesting that HyNkd is involved in the large-scale $\beta$-catenin-based patterning system; (d–f) formation of distinct patterns (diffusive vs. spotty) in animals treated with 5 $\mu M$ ALP, evaluated after 72 h, suggesting that expression levels of multiple *Wnts* are regulated differently from HyNkd.

### 6 S6: Knockdown validation

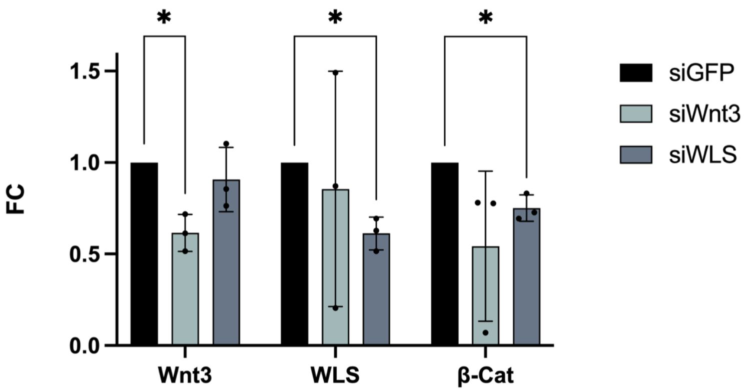


**Supplementary Figure 5.** qRT-PCR to validate knock-down efficiency of Wnt3 and Wntless siRNA. Experiment was performed in technical and biological triplicates. Bars represent mean with standard deviation. Analysis was performed using the ΔΔCt method. * p-value < 0.05
